# Atomistic modelling of lysophospholipids from the Campylobacter jejuni lipidome

**DOI:** 10.1101/2025.02.13.638095

**Authors:** Kahlan E. Newman, Astrid F. Brandner, Jonathan W. Essex, Syma Khalid

## Abstract

Lysophospholipids are an important class of lipids in both prokaryotic and eukaryotic organisms. These lipids typically constitute a very small proportion (<1%) of the bacterial lipidome, but can constitute 20-45% of the *Campylobacter jejuni* lipidome under stress conditions. It is thus of importance to include these lipids in model *C. jejuni* membrane simulations for an accurate representation of the lipidic complexity of these systems. Here, we present atomistic models for four lysophospholipids from the *C. jejuni* lipidome, each derived from existing phospholipid models. Herein we use molecular dynamics simulations to evaluate the ability of these models to reproduce the expected micellar, hexagonal, and lamellar phases at varying levels of hydration. Mixtures of phospholipids and lysophospholipids emulating the *C. jejuni* lipidome under ideal growth conditions were found to self-assemble into bilayers in solution. The properties of these mixed bilayers were compared to those containing only phospholipids: the presence of the selected lysophospholipids causes a subtle thinning of the bilayer and a reduction in area per lipid, but no significant change in lipid diffusion. We further test the mixed bilayer model running simulations in which a native inner membrane protein is embedded within the bilayer. Finally, we show that lysophospholipids facilitate the formation of pores in the membrane, with lysophospholipid-containing bilayers more susceptible to electroporation than those containing only phospholipids.

## Introduction

### Lysophospholipids: Synthesis & Function

Often the assumption when referring to phospholipids is a molecule with a hydrophilic headgroup connected to two hydrophobic acyl tails *via* a glycerol 3-phosphate group. However, phospholipids may have an additional acyl chain, or may have just one tail; such lipids are termed acylphospholipids and Lysophospholipids (LPLs), respectively. LPLs are intermediates in phospholipid synthesis^1^ and metabolites of phospholipid degradation^2–5^ (Fig. 1).

**Figure 1:**
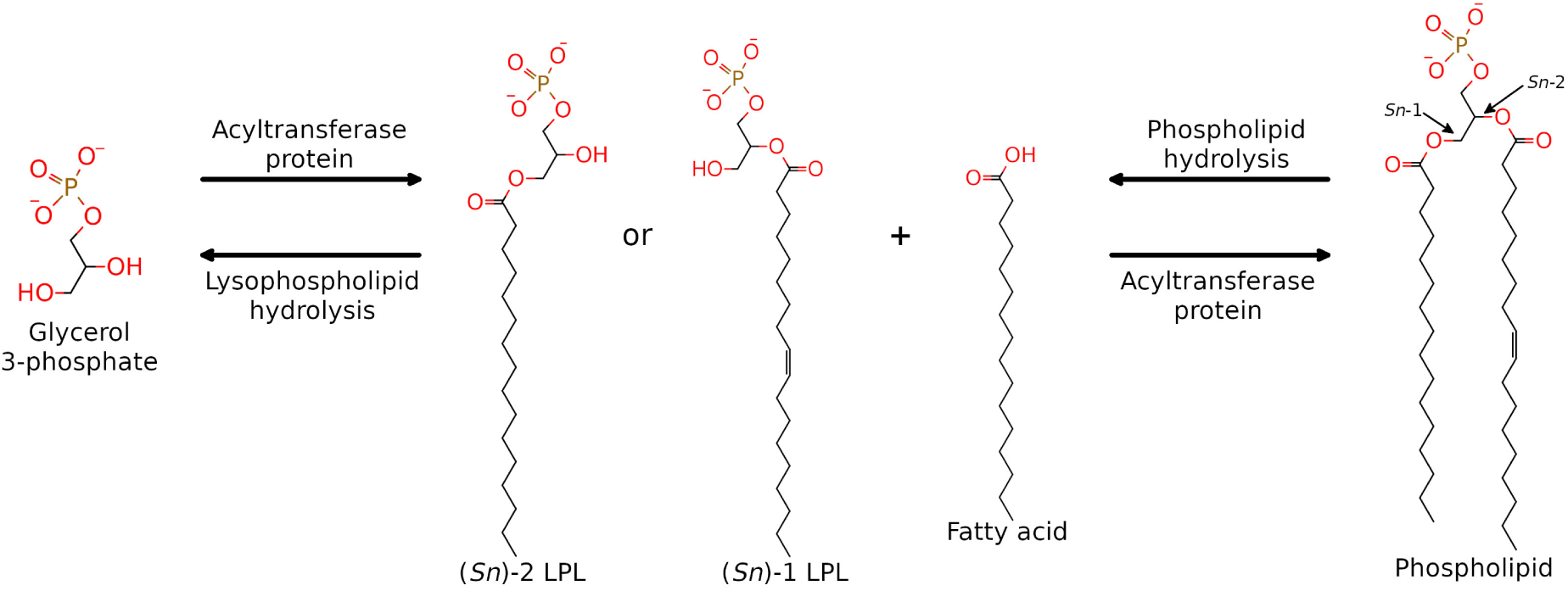
LPLs are generated as intermediates in phospholipid biosynthesis and as by-products of phospholipid degradation.

LPLs are known to be important in signalling, lipid raft formation, and membrane remodelling in eukaryotes. ^6^ The role(s) of LPLs in bacteria remain poorly understood, although they are thought to be involved in bacterial survival and invasion.^6,7^ Diverse mixtures of lipids likely aid bacteria in maintaining their shape, which in turn is important for function, *e.g.* the helical shapes of *Campylobacter* and *Helicobacter* are important for motility in the gut. ^8^ LPLs have been shown to constitute a substantial proportion of the *C. jejuni* lipidome. ^9^ Under ideal growth conditions the *C. jejuni* lipidome was shown to contain 45% Phosphatidylglycerol (PG), 28% Phosphatidylethanolamine (PE), 16% lysoPE, 4% PX (unknown headgroup), 3% lysoPG, 2% Phosphatidic acid (PA), and 1% acylPG, determined *via* high-performance liquid chromatography tandem-mass spectrometry. ^9^ Under stress conditions, the proportion of LPLs increased to as much as 45%. ^9^ This is in contrast to other bacterial species where LPLs are typically found in concentrations below 1%.^2^ This high lysophospholipid content may be due to the absence of the LplT-Aas ‘phospholipid repair’ system in *C. jejuni* : ^9^ LplT proteins translocate LPLs across the inner membrane of Gram negative bacteria where they can be re-acylated by the acyltransferase protein Aas on the cytoplasmic face of the membrane. ^2,10,11^ Without this system, LPLs can accumulate in the membranes of the bacterium. LPLs are thus of interest for modelling a biologically relevant *C. jejuni* membrane.

### Lipid Geometry

Phospholipid molecules can adopt a range of shapes based on their headgroup, number of acyl tails, tail length, and tail saturation. This in turn affects the assemblies these lipids form. The shape of a lipid can be described by its Critical Packing Parameter (CPP):^12^

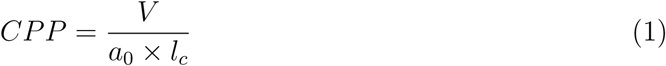

where *V* is the hydrophobic tail volume; *a*_0_ is the optimal headgroup area; and *l_c_* is the length of the hydrocarbon tail. *V* and *l_c_* can be derived analytically as they are dependent only on the number of carbon atoms in the hydrophobic tail(s). ^13^ Determination of *a*_0_, however, is non-trivial. This value is usually resolved experimentally, though different methods can yield disparate results. ^14,15^ Molecular Dynamics (MD) simulations of lipid monolayers have been used to quantify *a*_0_:^16^ the shapes of the lipid species were generally in agreement with the known geometries, but quantitative agreement between calculated and experimentally-determined values for *a*_0_ (and subsequently CPP) was limited.^16^

The geometry of individual lipid molecules, defined by the CPP, largely determines their resulting aggregates in aqueous solution (Fig. 2). LPLs have a single acyl tail: due to the reduced ratio of *V* to *a*_0_*l_c_* these lipids are of a conical geometry and typically self-assemble into micelles in solution. ^12,17–19^ Pure LPLs are non-bilayer-forming lipids under physiological conditions. ^20^ However, LPLs can stabilise membranes containing multiple lipid types *via* shape complementarity, *e.g.* in a bilayer rich in inverted conical lipids such as cardiolipin.^21^ Conversely, LPLs can disrupt membranes through detergent-like character, increasing the permeability of phospholipid bilayers. ^22–24^

**Figure 2:**
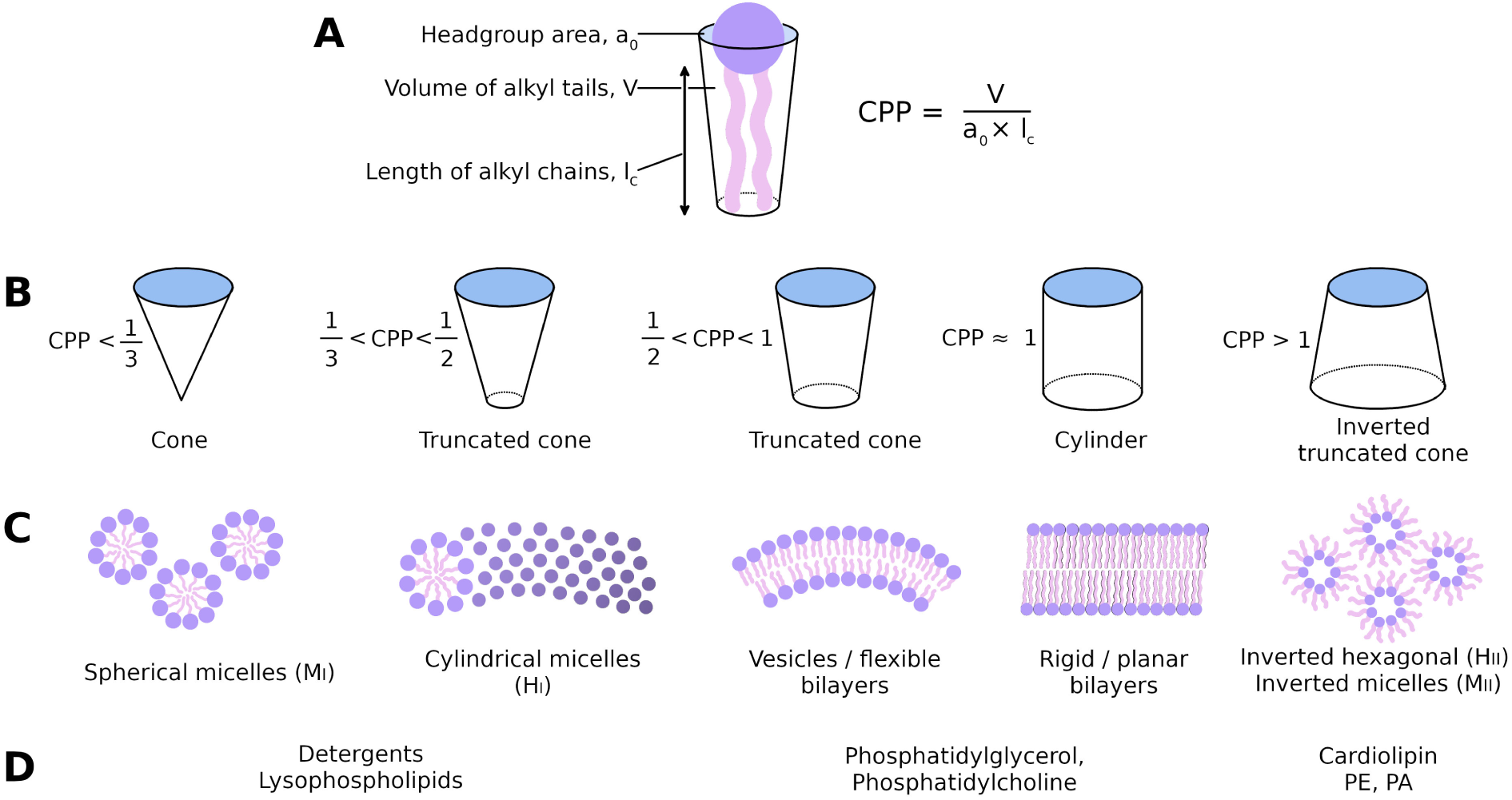
Lipid shapes and resulting aggregates. Adapted from Ref. 25. **(A)** Visualisation of the parameters defining the Critical Packing Parameter, CPP (Eq. 1). **(B)** Approximate thresholds for the value of CPP for geometry classification. ^26^ **(C)** Aggregates arising from lipids of different geometries. **(D)** Examples of lipids of the described geometries.

Assemblies can also be affected by pH, salt concentration, temperature, and hydration level. ^18,19,27^ Phase diagrams relating temperature and hydration have been resolved for some LPLs: ^19^ the phase diagrams for 1-palmitoyl-2-hydroxy-sn-glycero-3-phosphatidylcholine (lysoPC_(16:0)_) and 1-stearoyl-2-hydroxy-sn-glycero-3-phosphatidylcholine (lysoPC_(18:0)_) are shown in Fig. 3.^28^ In excess water LPLs typically self-assemble into micelles (M*_I_* phase). As the level of hydration decreases, different phases are known to form. For the lysoPC lipids shown in Fig. 3, hexagonal (H*_I_*), cubic (Q*_I_*), and lamellar (L*_α_*, L*_c_*) phases form as the ratio of water to lipids decreases. Similarly, at low temperatures lysoPE has been shown to achieve a metastable interdigitated lamellar gel phase.^17,18^

**Figure 3:**
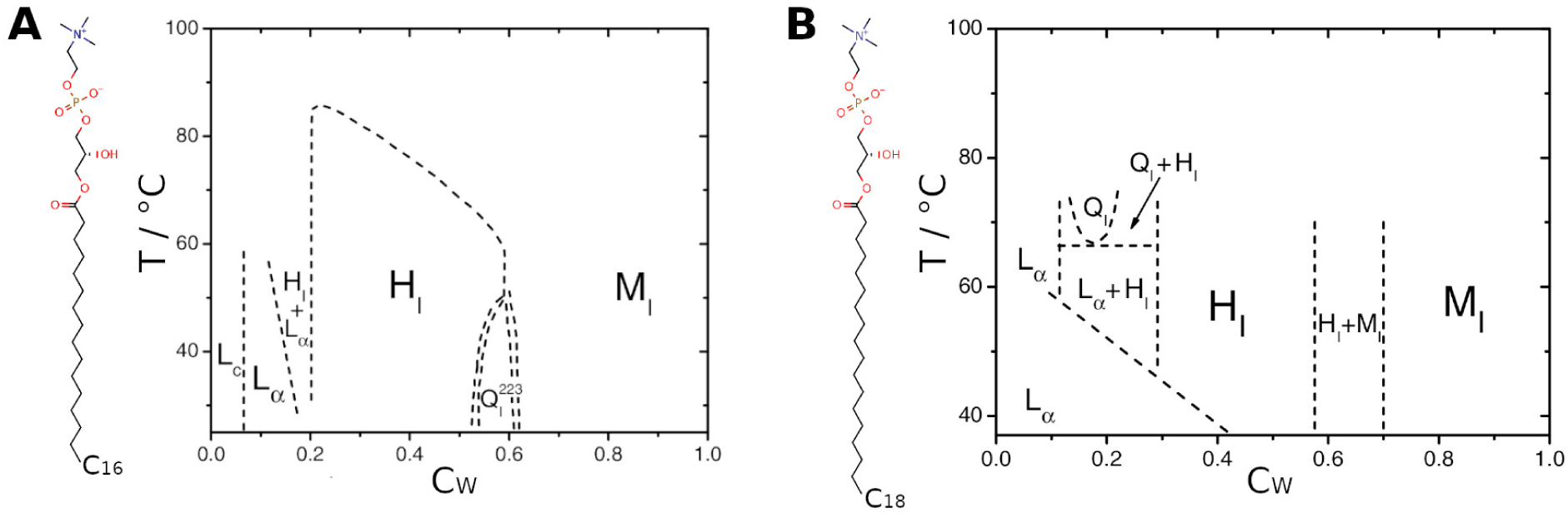
Phase diagrams for lysoPC lipids, ^18^ adapted from Ref. 19. L*_C_*, solid crystalline phase; L*_α_*, liquid crystalline (fluid) bilayer phase; 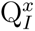, cubic phase; H*_I_*, hexagonal phase; M*_I_*, micellar phase. **(A)** Structure and phase diagram of 1-palmitoyl-2-hydroxy-sn-glycero-3-phosphatidylcholine (lysoPC, 16:0). **(B)** Structure and phase diagram of 1-stearoyl-2-hydroxy-sn-glycero-3-phosphatidylcholine (lysoPC, 18:0). Both of these LPLs aggregate into micelles in excess water, but form hexagonal, cubic, and lamellar phases when the water content (C*_W_*) is reduced.

### Lipid Modelling

Due to a lack of readily available models, LPLs were omitted in our previous simulation study of the *C. jejuni* outer membrane. ^29^ However, due to their wide-ranging effects on membranes and embedded proteins,^30,31^ as well as their high concentration in this bacterial lipidome, we feel it important to include this family of lipids in further *C. jejuni* membrane models. Four of the most abundant LPLs were selected from the *C. jejuni* lipidome^9^ for modelling: lysoPE (16:0, 18:1, and 19:0c) and lysoPG (18:1) (Fig. 4).

**Figure 4:**
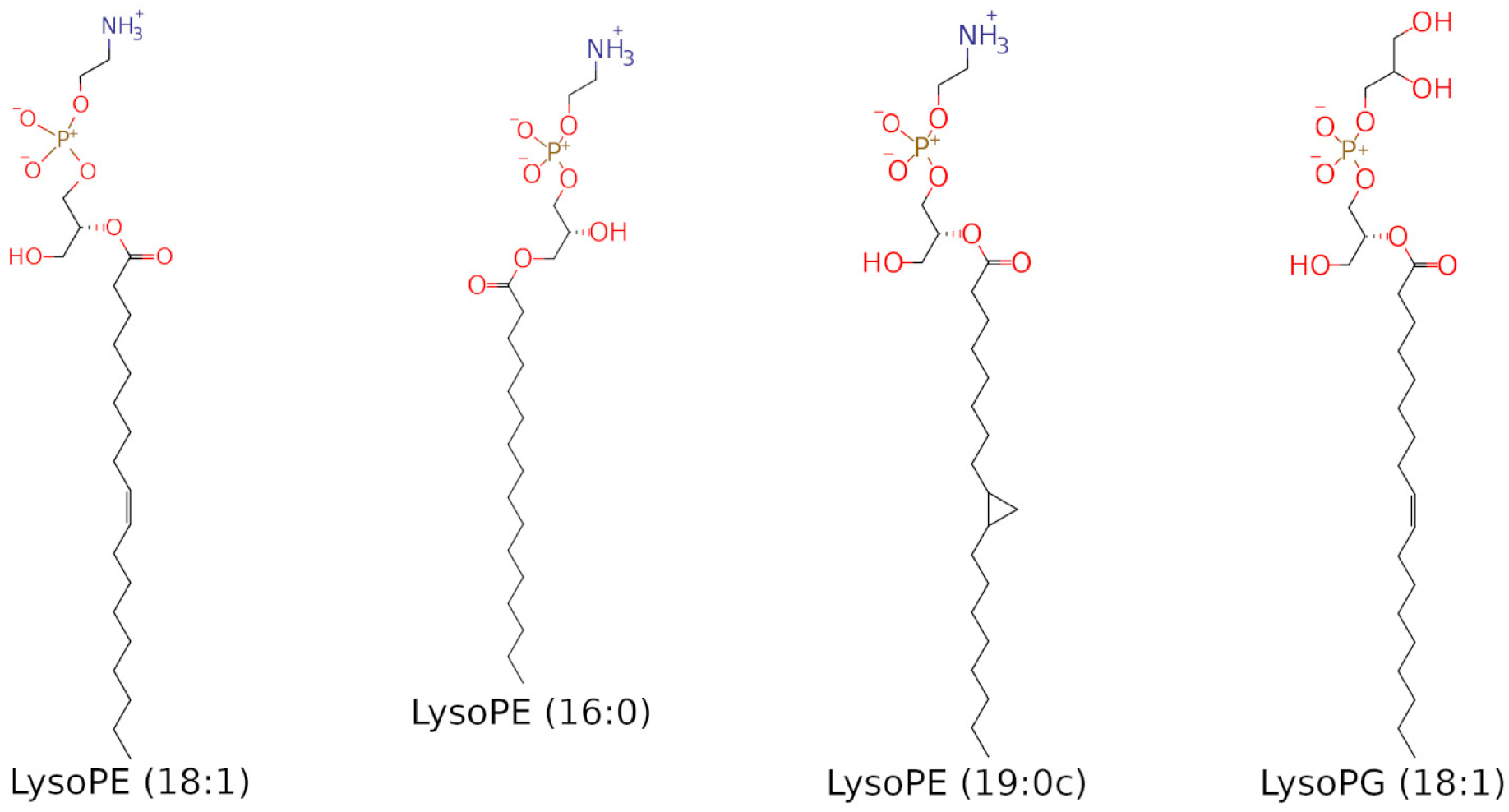
Structures of the lysophospholipids modelled. Three lysoPE lipids and one lysoPG lipid were selected from the *C. jejuni* lipidome.

The development of parameters for individual molecular models is ordinarily an iterative process, wherein an initial model is refined to better match experimental (or quantum mechanical) data.^32^ However, experimental data for all but the most common lipid species is sparse; to the best of our knowledge, there is no experimental data available to iteratively improve any models we generate specifically relating to the selected lipids. It is also important to note that data alone is insufficient for parameterisation of large biomolecules. ^33^ While *ab initio* data is useful to parameterise quantities such as dihedral energy surfaces and optimal bond lengths, it cannot accurately capture non-bonded parameters relevant to Lennard-Jones potentials and electrostatics, which are key for capturing phase behaviour. We will therefore qualitatively evaluate our models against general properties of lysophospholipids. To this end, we have produced models for four LPLs by modifying existing, experimentally-validated lipid models in the CHARMM36m force field, with the aim to generate more diverse bilayers that better represent the complexity of *C. jejuni* membranes. Here we have tested these models *via* aqueous self-assembly simulations and by evaluating their effects on the properties of phospholipid bilayers.

## Materials and Methods

All MD simulations were performed in the CHARMM36m forcefield^34,35^ with modified TIP3P water^36^ in GROMACS 2021.5. ^37,38^ Hydrogen to heavy atom bonds were constrained using LINCS.^39^ Long-range electrostatics were treated using Particle Mesh Ewald (PME)^40^ (1.2 *nm* cut-off). Van der Waals interactions were smoothed to zero between an inner cut-off of 1.0 *nm* to a final cut-off of 1.2 *nm*. Unless otherwise stated: the velocity rescale thermostat^41^ (*τ_T_* = 1.0 *ps*) was used to couple systems to a heatbath; pressure coupling used a time constant, *τ_P_*, of 2 *ps* and compressibility, *β*, of 4.5 *×* 10*^−^*^5^ bar*^−^*^1^; and a timestep of 2 *fs* was used throughout. Analyses were performed using GROMACS, MDAnalysis, and LiPyphilic utilities^42–46^ and in-house scripts. Molecular graphics were generated in VMD 1.9.4a55.^47^ Aggregates split across periodic boundaries were reassembled using FixBox.^48^

### Model Generation

Models were derived from existing phospholipid models in the CHARMM36m forcefield. ^49,50^ LysoPE lipids were generated by modifying 1-palmitoyl-2-oleoyl-sn-glycero-3-phosphoethanolamine (POPE) and 1-palmytoil-2-cis-9,10-methylenehexadecanoyl-phosphatidylethanolamine (PMPE) models; lysoPG was generated by modifying the 1-palmitoyl-2-oleoyl-sn-glycero-3-phosphoglycerol (POPG) model. In each case, the relevant lipid tail was removed, and the linking ester group replaced with a hydroxyl group. Parameters for the O-H bond and hydroxyl hydrogen were taken from the CHARMM36m model for the lysophospholipid 1-myristoyl-2-hydroxy-sn-glycero-3-phospho-(1’-rac-glycerol) (LMPG)(saturated C_14_ tail). Where necessary, the partial charge of the headgroup phosphorus atom was modified (at most *±* 0.08 *e*) to yield the correct net integer charge. The resulting models were energy minimised *in vacuo* in 5,000 steps using steepest descent.^51^

### Self-Assembly Simulations

How the LPLs assemble in solution was assessed by simulating the lipids in water boxes. Lipid molecules were placed randomly in a simulation box using the GROMACS utility insert-molecules, with neutralising counterions (K^+^) plus additional KCl in solution (50-150 mM, system dependent). Several setups were explored, detailed in the following subsections. The contents and initial unit cell sizes of all simulated systems are detailed in the Supplementary Materials (Tables A1 to A7). All self-assembly simulation cells were initially cubic.

### Small Single-Lysophospholipid Systems

In this setup, each box contained 140 lysophospholipid molecules, plus ions and water molecules such that the ratio of lipids to water was either 1:50 (Water weight fraction (C*_W_*) ≈ 0.65-0.7), C*_W_* = 0.4, or C*_W_* = 0.1. Three unique box configurations were generated for each lipid at each concentration. Systems were energy minimised in 5,000 steps (steepest descent^51^). This was followed by NVT and NPT equilibration stages. In the NVT stage, the system was coupled to a heat bath at 315 *K* for 100 *ps*, with position restraints of 20 *kJmol^−^*^1^*nm^−^*^2^ on all lipid heavy atoms. The NPT stage introduced pressure coupling *via* the isotropic Parrinello-Rahman barostat^52,53^ to bring the system to a target pressure of 1 *bar* over 100 ps. Temperature coupling was maintained during the NPT stage. Each system was then simulated at 315 *K* under the isotropic Parinello-Rahman barostat at 1 bar. Simulations lasted 1 *µs* for excess water (C*_W_* = 0.65-0.7), and 1.5 *µs* for C*_W_* of 0.4 or 0.1. One replicate for each lipid species at C*_W_* 0.1, 0.4 was extended to 2 *µs*.

#### Larger Single-Lipid Systems

Larger systems containing more lipids were also simulated. For each lysophospholipid, three unique configurations were generated containing: 500 lipid molecules; water such that C*_W_* = 0.7, 0.4, or 0.1; neutralising counterions and KCl to 80-150 *mM* (system dependent). Systems were energy minimised and equilibrated as above. To allow the volume and density of the box to stabilise, a 5 *ns* unrestrained simulation under the conditions as described for the small single-lysophospholipid systems (isotropic pressure coupling) was applied after the equilibration stages. Each system was then simulated with anisotropic pressure coupling at 1 *bar* (Parrinello-Rahman). Simulated annealing was applied to discourage ‘trapping’ of the systems in metastable phases: each system was simulated at 315 *K* for 40 *ns* followed by 10 *ns* at 400 *K*, cycling for 500 *ns* in total.

#### Small Mixed-Lipid Systems

To assess whether a bilayer containing both phospholipids and LPLs would form spontaneously, we repeated the above isotropic simulations (small single-lysophospholipid systems) with phospholipids and phospholipid-lysophospholipid mixtures. Three unique systems were established for each pure phospholipid species: 140 POPE, 1-palmitoyl-2-oleoyl-sn-glycero-3- phosphatidic acid (POPA), or POPE molecules were each placed randomly in boxes with 50 water molecules per lipid, counterions and KCl to 80 *mM*. Three unique boxes containing a lipid mixture with 20% LPLs were also generated: molecules of POPG, POPE, POPA, and the four LPLs were placed in the simulation box in a 9:6:1:1:1:1:1 ratio. These systems were energy minimised, equilibrated, and simulated for 1 *µs* under the conditions described for the small single-lysophospholipid systems.

### Equilibrium Bilayer Simulations

The self-assembled 20% LPLs bilayer systems were tiled in a 3*×*3 grid to yield larger bilayers. The three replicates were combined in three unique configurations, reflected in the *xy* plane where necessary to minimise asymmetry with a small gap between tiles to avoid steric clashes. After energy minimisation (5,000 steps steepest descent), these gaps in the bilayer were closed using 2*×*1 *ns* NPT stages at 300 *K*. Position restraints were applied to the *z* coordinates of lipid heavy atoms to allow only lateral translation. Systems were coupled to heat and pressure baths using the Berendsen regime^54^ in both stages. A semi-isotropic barostat was employed; a pressure of 1.2 *bar* was applied in the *xy* plane (*β* = 4.5 *×* 10*^−^*^3^ *bar^−^*^1^) to encourage compression, and a pressure of 1 *bar* was applied along the *z* axis (*β* = 4.5 *×* 10*^−^*^5^ *bar^−^*^1^). Position restraints (500 *kJmol^−^*^1^ *nm^−^*^2^) were applied to lipid P and C/N/O atoms in the first nanosecond, and restraints of 200 and 50 *kJmol^−^*^1^*nm^−^*^2^, respectively, were applied in the second nanosecond. Each bilayer was then solvated in 150 *mM* KCl with additional neutralising counterions. Three analogous phospholipid-only bilayers (POPG, POPE, POPA, 9:6:1) of comparable size were generated using CHARMM-GUI Membrane Builder. ^49,55,56^

Systems were energy minimised in 50,000 steps (steepest descent), then equilibrated sequentially using 2 NVT and 4 NPT stages, decreasing position restraint strength at each stage (Table 1). All NPT stages coupled the system to a pressure bath at 1 *bar* using the semi-isotropic Berendsen scheme^54^ (*τ_p_* = 5.0 *ps*). This was followed by 1 *µs* unrestrained production simulation. Systems were coupled to a temperature bath of 315 *K* using the Nosé-Hoover thermostat (*τ_T_*= 1.0 *ps*) and a pressure bath of 1 *bar* using the semi-isotropic Parrinello-Rahman regime (*τ_P_* = 5.0 *ps*).

**Table 1:** Restraints for lipid phosphorus atoms and dihedrals; timestep (dt); and duration for equilibration stages.

| Equilibration stage | Position restraint / $\text{kJmol}^{-1} \text{ nm}^{-2}$ | | dt / fs | Length / ps |
| --- | --- | --- | --- | --- |
|  | Lipid P | Dihedrals |  |  |
| NVT1 | 1000 | 1000 | 1 | 125 |
| NVT2 | 400 | 400 | 1 | 125 |
| NPT1 | 400 | 200 | 2 | 500 |
| NPT2 | 200 | 200 | 2 | 500 |
| NPT3 | 40 | 100 | 2 | 500 |
| NPT4 | - | - | 2 | 500 |

### Native Protein embedded in Bilayer System

SWISS-MODEL^57^ was used to fit the PglB sequence of *C. Jejuni* to the *C. lari* PglB structure (PDB ID: 5OGL^58^). Default protonation states were assigned for all residues in this model. PglB was glycosylated using the CHARMM-GUI PDB Reader and Manipulator module^59–61^ with the following glycan sequence:

GalNAc-*α*1,4-GalNAc-*α*1,4-[Glc-*β*1,3]GalNAc-*α*1,4-GalNAc-*α*1,4-GalNAc-*α*1,3-diNAcBac-*β*1

where diNAcBac is N*^′^*,N*^′^*-diacetylbacillosamine [2,4-diacetamido-2,4,6 trideoxyglucopyranose]; GalNAc is N-acetylgalactosamine; and Glc is glucose. This template contained two magnesium ions, a donor substrate (Lipid-Linked Oligosaccharide (LLO), with a truncated lipid tail), and an acceptor peptide. The magnesium ions were maintained at the locations from the template. The structure of the acceptor peptide for PglB was extracted from the X-ray structure 5OGL. ^58^ The sequence was modified to match that of the known occupied CmeB glycosite (KDRNVSAD) using *in silico* mutations in PyMOL [488]. The glycan donor LLO was generated in CHARMM-GUI Ligand Modeler^62^ with the lipid tail of the LLO truncated to the same length as that in the PglB crystal structure. The lipid tail and bacillosamine unit were aligned with those of the crystal structure. The remaining 6 moieties of the heptasaccharide were manually arranged. This substrate- bound protein was energy minimised (5,000 steps steepest descent^51^) to resolve any steric clashes and inappropriate bond lengths/angles/dihedrals that had arisen. The complex containing PglB was embedded in a membrane patch obtained from a previously equilibrated 20% LPL mixed bilayer, so as to match the hydrophobic trans-membrane domains of the protein with the bilayer core. The full system was energy minimised *via* steepest descent algorithm for 50,000 steps, equilibrated in NPT with a 1 *fs* timestep for 100 *ps* using a Berendsen barostat with semi-isotropic coupling and coupling (1 bar, *τ_P_* = 5.0 *ps*), the temperature *via* a Berendsen thermostat (*τ_T_* = 1.0 *ps*, reference temperature of 315 *K*). Three replicas were subjected to the production run protocol, firstly for 1 *ns* only, and subsequently each of the replicas was run for 1 *µs* for analysis with a 2 *fs* integration timestep. The full details of these system are available in Supplementary Materials, Table A6. In the production run, systems were coupled to a temperature bath of 315 *K* using the V-rescale thermostat (*τ_T_*= 1.0 *ps*) and a pressure bath of 1 *bar* using the semi-isotropic Parrinello-Rahman regime (*τ_P_* = 5.0 *ps*, compressibility = 4.5e*^−^*^5^ *bar^−^*^1^). GROMACS rms and rmsf tools, MDAnalysis^45^ and fatslim^63^ were used to create in house scripts for data analysis. Phosphorus atoms were used to define the headgroups for thickness and area per lipid analysis. A contact between sugar and lipid was computed if the distance between the sugar moieties of the LLO donor lipid and any lipid atom of the corresponding lipid type was within 0.4 *nm*. The contacts were only counted once per glycan-lipid interaction, regardless of the total number of atoms in contact. When mean values and their associated errors are reported, unless otherwise stated, values were computed for each replica based on the last 500 *ns* of production run.

### Electroporation Simulations

Three mixed phospholipid-LPL bilayers were generated as described in the previous section using a 2*×*2 grid (4 tiles total). In this case, each bilayer was generated from 4 copies of a single self-assembled bilayer replicate; two tiles were flipped in the *xy* plane to generate fully symmetric bilayers. Three bilayers of comparable size containing only phospholipids (9:6:1 POPG, POPE, POPA) were generated in CHARMM-GUI Membrane Builder. Bilayers were solvated in 150 *mM* KCl, minimised, equilibrated, and subjected to 100 *ns* unrestrained equilibrium simulation as described for the 3*×*3 tiled bilayers. An electric field was then applied to each system along the *z*-axis. Field strengths of 0.100, 0.125, 0.150, 0.175, 0.200 *V nm^−^*^1^ were applied, and the time taken for each bilayer to electroporate was measured.

Each of the porated bilayers generated in the 0.200 *V nm^−^*^1^ simulations (3*×* phospholipid only, 3 *×* 20% LPLs) were simulated under equilibrium conditions to assess whether the pores would close in the absence of an external electric field. Initial structures for these subsequent simulations were taken as the frame at which the box *x* dimension had increased to 110% of its equilibrium value. These systems were simulated for 100 *ns* under the conditions described for the equilibrium bilayer simulations.

## Results and discussion

### Micelle Formation in Excess Water

LPLs in excess water are known to form micelles (Fig. 3). ^18,19^ Our simulations of these four lysophospholipids are consistent with this observation. Each lipid model was simulated in solution at C*_W_* = 0.65-0.7 in a small box (140 lipids) under isotropic pressure coupling for 1 *µs*. Approximately spherical micelles formed in most cases (Fig. 5A). At the start of most production simulations, multiple micelles form as the hydrophobic tails aggregate; usually this was two micelles, each containing approximately half of the lipids (Fig. 5B). As the simulations progress, these aggregates combine to form a single micelle containing all 140 LPLs that is roughly spherical (Fig. 5C). In two simulations, a single micelle containing all the lipids formed immediately. In eight of the twelve simulations, these spherical micelles were maintained for the remainder of each simulation.

**Figure 5:**
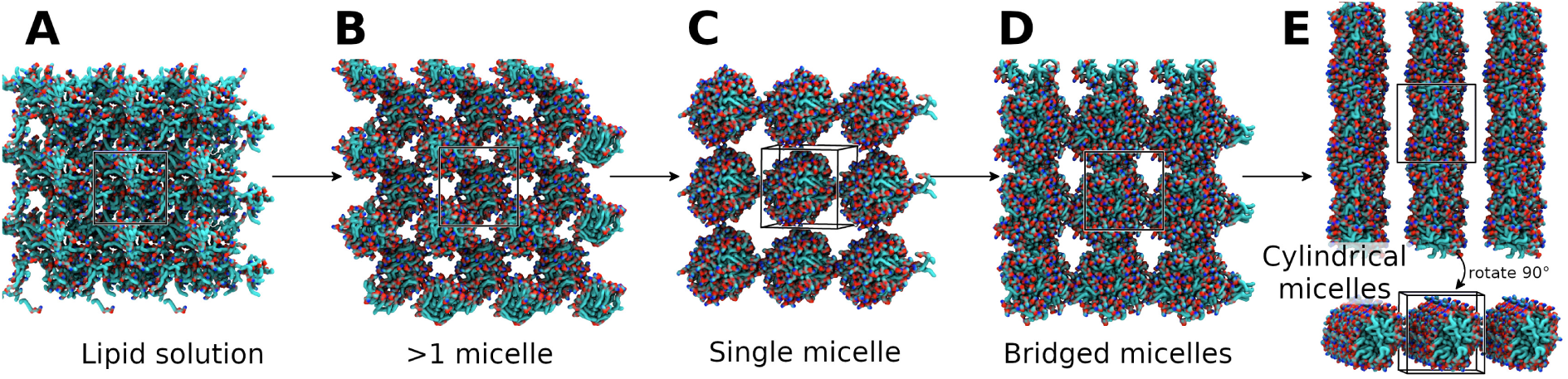
Formation of micelles in solution. Snapshots from lysoPE_(19:0_*_c_*_)_ R1, 50 waters per lipid, isotropic pressure coupling. Lipids shown in surface representation, coloured by element. Unit cell shown with black lines. Water and ions omitted for clarity. **(A)** Initial lipid solution. **(B)** Lipid tails aggregate to form multiple (2) smaller micelles. **(C)** Smaller micelles combine to form a single spherical micelle. **(D)** Lipid bridges form between one side of the micelle and the opposite side in the adjacent periodic image. **(E)** Bridged micelles rearrange to form cylindrical micelles that are continuous across periodic boundaries in some replicates.

In 3*×*lysoPE_(19:0_*_c_*_)_ replicates and a single lysoPE_(18:1)_ replicate, where the shape of the lysophospholipid was less conical, cylindrical micelles formed. These simulations also initially proceed *via* spherical micelle aggregates (Fig. 5A-C). One side of the micelle then interacts with the other through its periodic image, allowing lipid bridges to form; a single cylindrical micelle forms over periodic boundaries (Fig. 5D,E).

Further simulations of these lipids in solution in a larger box (500 lipids, C*_W_* = 0.7) with anisotropic pressure coupling displayed similar micelle formation. In each case, multiple micelles formed in each simulation box. However, as there were no extended, rigid structures formed over the periodic boundaries, there is a lack of both cohesive forces that would prevent expansion and of repulsive forces that prevent collapse of the simulation cell along a given axis. Box deformation and subsequent collapse (a box dimension became smaller than the electrostatic cut-off) occurred rapidly in all replicates (Supplementary Materials, Fig. A1).

### Aggregates at Higher Lipid Concentrations

#### Small Isotropic Boxes

LPLs at a higher concentration/lower C*_W_* self-assemble into other phases, such as hexagonal, cubic, and lamellar phases^18,19^ (Fig. 3). Each lysophospholipid model was simulated in solution at C*_W_* = 0.4 and 0.1 in a small box (140 lipids) under isotropic pressure coupling for 1.5-2 *µs* to assess whether these phases would also form. These simulations displayed some success in reproducing the expected phases. At C*_W_* = 0.1, four of the twelve replicates approach a lamellar phase, with one replicate of lysoPE_(19:0_*_c_*_)_ forming a multilayer L*_α_* system in 1.25 *µs* (Fig. 6A,B). One replicate each of lysoPE_(16:0)_, lysoPE_(18:1)_, and lysoPG_(18:1)_ also approach a multilayered lamellar phase, but do not reach the L*_c/α_* phase on the timescales simulated; while a clearly layered aggregate is formed, headgroups remain in what should be an exclusively hydrophobic region (Fig. 6A). As this arrangement is an intermediate stage of the bilayer formation for the lysoPE_(19:0_*_c_*_)_ replicate, it is likely that a lamellar phase may form if these replicates were extended.

**Figure 6:**
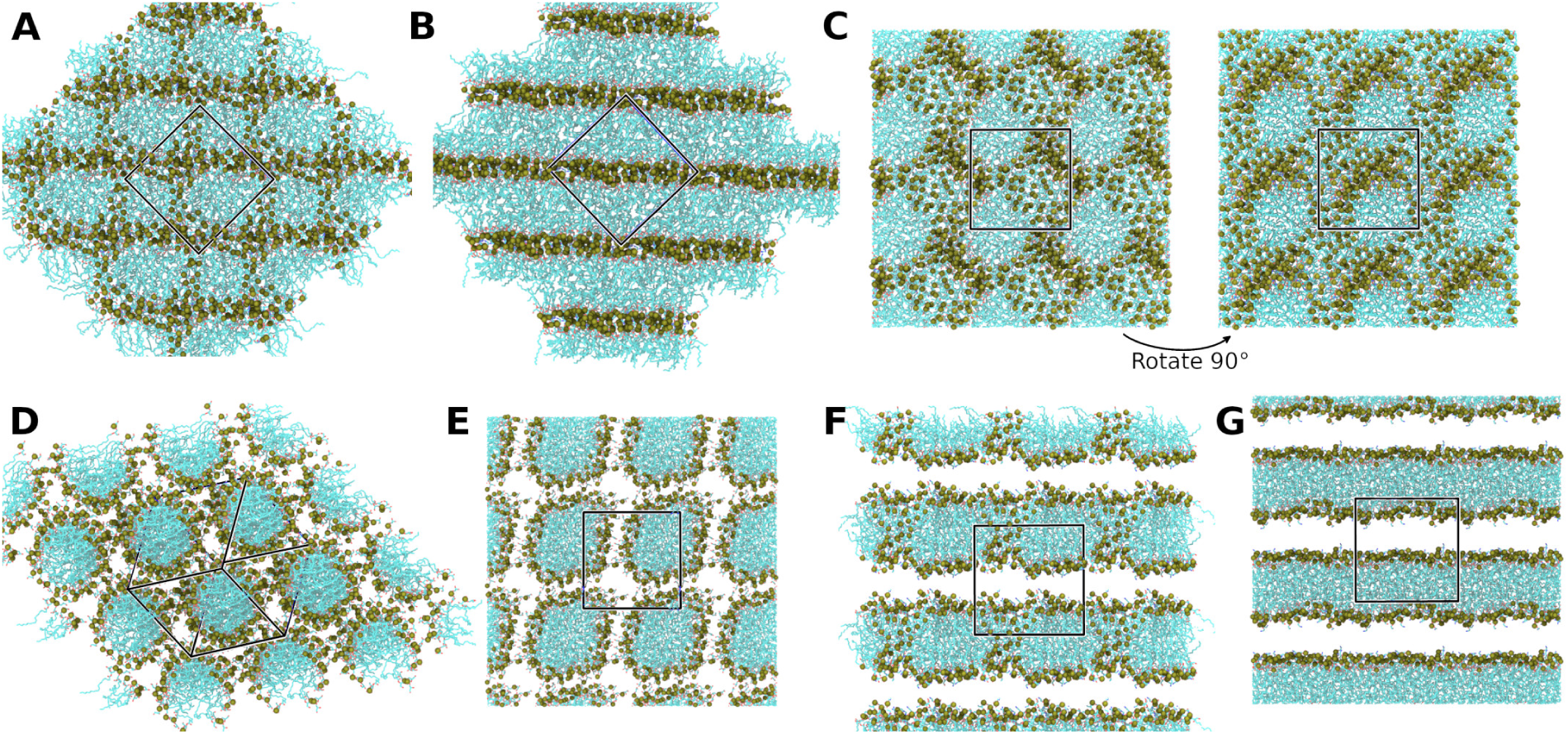
Final phases formed by the modelled lysophospholipids at low hydration under isotropic pressure coupling. **(A-C)** snapshots from C*_W_* =0.1 systems. **(D-G)** Snapshots from C*_W_* =0.4 systems. Unit cells indicated by a black box. Lipids coloured by element in stick representation, with headgroup phosphorus atoms shown as tan spheres. Water and ions omitted for clarity. **(A)** LysoPE_(18:1)_ approaching a multilayered lamellar phase; there are still headgroups within the hydrophobic core of the bilayers. One replicate each of lysoPE_(16:0)_ and lysoPG_(18:1)_ also reached this phase. **(B)** Multilayered L*_α_* phase formed by a single replicate of lysoPE_(19:0_*_c_*_)_. **(C)** Cubic phase formed in eight of the twelve simulations at C*_W_* = 0.1. **(D)** Hexagonal (H*_I_*) phase formed by a single replicate of lysoPG_(18:1)_. **(E)** Primitive cubic arrangement of cylindrical micelles, formed by three replicates of lysoPE_(16:0)_ and two replicates of lysoPG_(18:1)_. **(F)** A porated bilayer formed by all three replicates of lysoPE_(18:1)_.**(G)** L*_α_* phase formed by a single replicate of lysoPE_(19:0_*_c_*_)_.

The other eight replicates at C*_W_* =0.1 formed a state that resembled neither a lamellar nor a hexagonal phase. Instead these replicates displayed cubic phases, wherein highly interconnected micelle-like aggregates organise in a primitive-cubic unit (Fig. 6C). Where this state formed, it was stable for the remainder of the simulation.

At C*_W_* = 0.4 the aggregates formed can be clustered into two broad categories: cylindrical micelle-based assemblies; and bilayer-like assemblies. In six of the twelve systems, cylindrical micelles formed: in one replicate (1*×*lysoPG_(18:1)_) a hexagonal arrangement of these aggregates is observed, consistent with the expected H*_I_* phase (Fig. 6D). In the other five replicates (3*×*lysoPE_(16:0)_; 2*×*lysoPG_(18:1)_) these were in a primitive cubic arrangement (Fig. 6E). The remaining six replicates formed bilayer-like aggregates. All three replicates of lysoPE_(18:1)_ and two replicates of lysoPE_(19:0_*_c_*_)_ were observed to assemble into a porated bilayer (Fig. 6F). The final replicate of lysoPE_(19:0_*_c_*_)_ was observed to enter the L*_α_* phase (Fig. 6G) within the first 200 *ns* of the simulation. Longer simulations of these systems, than those feasible in the present study, would likely reduce these differences.

The use of isotropic pressure coupling and a small simulation box affects the likelihood of particular phases forming. When the box is small, the requirement for periodic replication can trap the system in a particular phase. Isotropic pressure coupling enforces uniform scaling along all box dimensions; the ability of a given lipid to form a lamellar bilayer will be dependent on selecting an appropriate box size and hydration level, or require a multilayered bilayer to form to appropriately span the periodic boundaries. Furthermore, while the simulations were relatively long in an attempt to sample more of the available phase space, most replicates became trapped in the first stable/metastable state they sampled. We attempted to address these issues with additional simulations under a different regime.

#### Larger Anisotropic Boxes with Simulated Annealing

To further assess the formation of these phases, each lipid model was simulated in solution at C*_W_* = 0.4 and 0.1 in a larger system (500 lipids) under anisotropic pressure coupling with simulated annealing. The use of anisotropic pressure coupling allows more degrees of freedom for extended structure formation and reduces templating effects as the simulation cell can deform in each dimension independently. Simulated annealing enhances sampling by temporarily increasing the kinetic energy of the system, encouraging the crossing of energy barriers and reducing the risk of the system becoming trapped in a metastable state.

At C*_W_* = 0.1, four of the twelve simulations approached a multilamellar assembly (Fig. 7A): two replicates each lysoPE_(18:1)_ and lysoPE_(19:0_*_c_*_)_ reach this state by the end of the 500 *ns* simulations. A fully lamellar assembly (where all headgroups have retreated from the hydrophobic core) was not achieved by any replicate on the timescale simulated. All other replicates at C*_W_* = 0.1 formed cubic assemblies as observed in the smaller isotropic systems (Fig. 6C).

**Figure 7:**
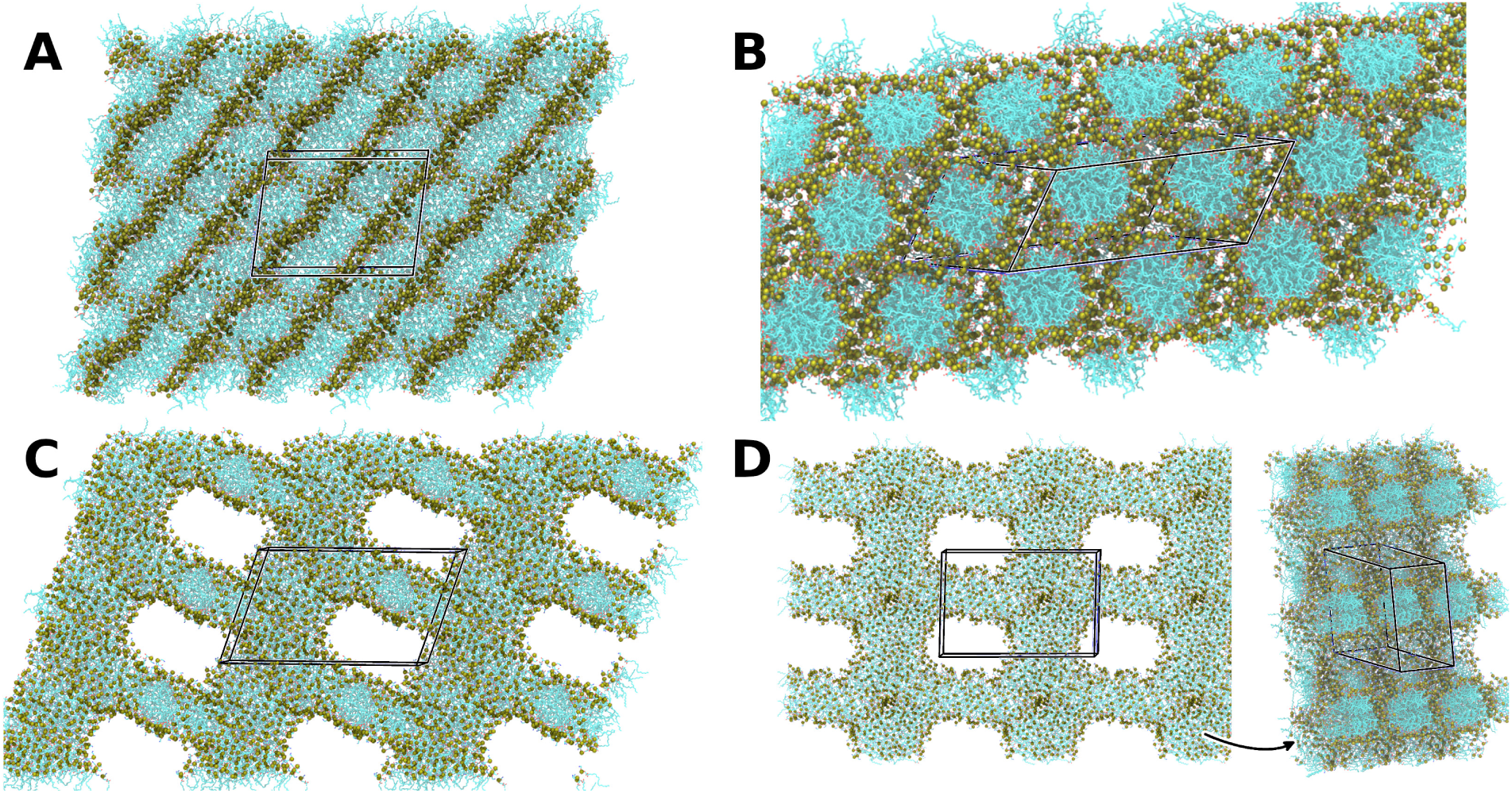
Final phases formed by the modelled lipids at low hydration under anisotropic pressure coupling. Unit cells indicated by a black box. Lipids coloured by element in stick representation, with headgroup phosphorus atoms shown as tan spheres. Water and ions omitted for clarity. **(A)** C*_W_* = 0.1. LysoPE_(18:1)_ approaching a multilayered lamellar phase; there are still headgroups within the hydrophobic core of the bilayers. Two replicates each of lysoPE_(18:1)_ and lysoPE_(19:0_*_c_*_)_ reached this phase. **(B-D)** C*_W_* = 0.4. **(B)** Hexagonal phase (H*_I_*) formed by 2 replicates of lysoPG_(18:1)_. **(C)** Mixed phase: combination of a bilayer-like assembly and a cylindrical assembly. **(D)** Cylindrical micelles in 2 directions, with lipid bridges at intersections.

The C*_W_* = 0.4 simulations displayed some diversity in the phases formed. Importantly, two replicates (2*×*lysoPG_(18:1)_) achieved the expected H*_I_* phase (Fig. 7B). The remaining replicates formed extended aggregates in two dimensions. In some cases, this was in the form of bilayer-like assemblies that extended across periodic boundaries in one plane, intersected by cylindrical micelles that extend across periodic boundaries along the perpendicular axis (Fig. 7C). In others, cylindrical micelles formed along two axes which intersect through lipid bridges (Fig. 7D). The simulation box collapsed within the first 300 *ns* in all replicates except those achieving the H*_I_* phase.

### Mixed Bilayer Formation

With the aim of simulating a bilayer more representative of the *C. jejuni* lipidome, we moved to assemble bilayers containing these LPLs. The *C. jejuni* lipidome (under ideal growth conditions) contains ∼20% LPLs: ^9^ we hypothesised that a mixture similar to that described in the literature would self-assemble into a bilayer. We first simulated the phospholipids POPG, POPE, and POPA in excess water (50 waters per lipid) to verify that these existing models were bilayer-forming.

Systems containing only POPA, POPE, or POPG in solution all formed bilayers within 400 *ns* under the simulated conditions. The mechanism of self-assembly was as described by Skjevik *et al.*^64^ (Fig. 8):

1. Initial solution of lipids, ions, and water.
2. Tails aggregate to form a micelle-like assembly. Lipid bridges form between one side of the micelle and the periodic image.
3. Bridging lipids insert into the assembly, forming a porous lamellar bilayer.
4. Lipid headgroups retreat from the hydrophobic core to the water-lipid interface; a non-porous bilayer is formed.

**Figure 8:**
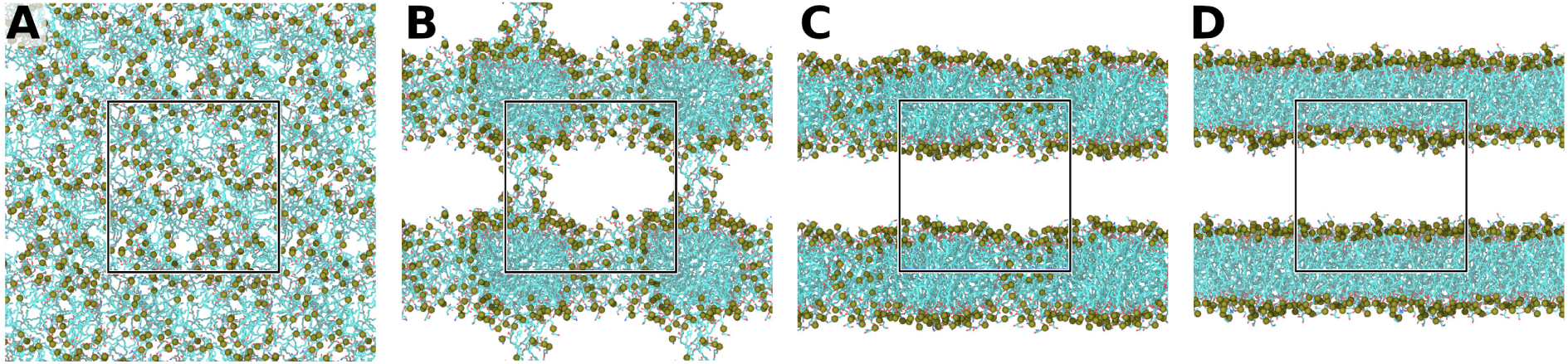
Stages of bilayer self-assembly as described in Ref. 64. Snapshots taken from self-assembly simulation of 20% LPL-phospholipid mixture. Lipids shown in stick representation coloured by element. Phosphorus atoms of the lipid headgroups highlighted as tan spheres. Water and ions omitted for clarity. Unit cell indicated by a black box. **(A)** Initial lipid solution. **(B)** Micelle-like assembly with lipid bridges between micelles and their periodic images. **(C)** Porated bilayer with lipid headgroups within the hydrophobic core region. **(D)** Non-porous bilayer formed.

These bilayers were stable for the remainder of the simulation time (1 *µs*). PE and PA lipids are of an inverted truncated cone shape, and may therefore be expected to form a more thermodynamically-favourable inverse hexagonal phase (H*_II_*). However, the energetic cost is sufficiently high to prevent the formation of non-lamellar phases under the simulated conditions. Each leaflet may tend to display negative curvature, but the leaflets stay together to maintain the hydrophobic core, resulting in a ‘frustrated bilayer’ ^65–67^

We then simulated a phospholipid-lysophospholipid mixture mimicking the proportions found in *C. jejuni* under ideal growth conditions. ^9^ Three systems, each containing 140 lipid molecules, were generated, with 20% lysophospholipid content: 63 POPG (45%); 42 POPE (30%); 7 POPA (5%); 7 lysoPE_(18:1)_ (5%); 7 lysoPE_(16:0)_ (5%); 7 lysoPE_(19:0_*_c_*_)_ (5%); 7 lysoPG_(18:1)_ (5%). Similar to the phospholipid-only systems, each replicate yielded a bilayer. The mechanism of bilayer self-assembly closely followed that described above (Fig. 8). A non-porous bilayer was formed in under 150 *ns* for 2 replicas and in ∼300 *ns* in the third. In all cases, these bilayers remained stable and non-porous for the remainder of the simulation (1 *µs*).

The contents of each leaflet were calculated to assess the level of asymmetry in the self-assembled bilayers (Table 2). While a strictly symmetric bilayer was not possible owing to an odd number of molecules for most species, the lysophospholipid content of each leaflet was considered. We would expect that the contents of each leaflet should approximately reflect the contents of the whole system, i.e. ∼20% LPLs, to minimise asymmetry and therefore curvature stress. For replicates 1 and 3 this was found to be the case, with the lysophospholipid content of each leaflet within 2.1 percentage points of the expected value. However, replicate 2 displayed greater asymmetry, with substantially more LPLs in the lower leaflet (18 *versus* 10).

**Table 2:** Contents of each leaflet in the self-assembled 20% LPL bilayers.

|  | R1 |  | R2 |  | R3 |  |
| --- | --- | --- | --- | --- | --- | --- |
|  | Upper | Lower | Upper | Lower | Upper | Lower |
| **POPG** | 32 | 31 | 33 | 30 | 31 | 32 |
| **POPE** | 24 | 18 | 21 | 21 | 18 | 24 |
| **POPA** | 3 | 4 | 5 | 2 | 4 | 3 |
| **LysoPE**<sub>(18:1)</sub> | 2 | 5 | 3 | 4 | 4 | 3 |
| **LysoPE**<sub>(16:0)</sub> | 4 | 3 | 4 | 3 | 4 | 3 |
| **LysoPE**<sub>(19:0c)</sub> | 3 | 4 | 1 | 6 | 2 | 5 |
| **LysoPG**<sub>(18:1)</sub> | 4 | 3 | 2 | 5 | 5 | 2 |
| **% LPL** | 18.1 | 22.1 | 14.5 | 25.4 | 22.1 | 18.1 |

Formation of the porated bilayers (Fig. 8C) occurred within 40 *ns* in each case. As lipid diffusion is relatively slow, the initial aggregation of the lipids will influence the final contents of each leaflet; once a stable/metastable bilayer arrangement is achieved, it is unlikely that this will change substantially without additional energetic input. Further to this, no lipid flip-flopping was observed during the simulations; the lipid content of each leaflet was constant once the bilayer had formed in each case. This is expected as the energy barrier to lipid flip-flop events is large (typically *>*20 *kJmol^−^*^1^ ^68,69^) resulting in timescales on the order of hours to weeks. ^70,71^

We did not observe any large persistent clusters of LPLs within the formed bilayers. While the lateral distribution of the LPLs in the self-assembled bilayers was not uniform, there was no obvious phase separation or clear enrichment of LPLs that was maintained for long periods of time (Fig. 9; Supplementary Materials, Table A8). However, the stochastic dimerisation of LPLs over the leaflets does appear to be important in poration events; this is discussed in the Electroporation section.

**Figure 9:**
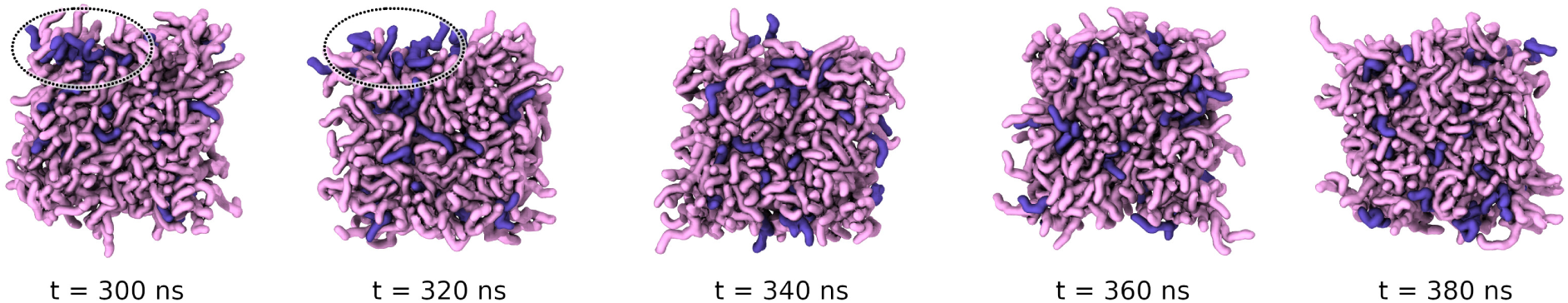
Lateral distribution of lysophospholipids in a self-assembled bilayer. Shapshots from replicate 1 of the self-assembled bilayer at different times. Lysophospholipids shown in purple, phospholipids in pink. While there is occasionally some aggregation of the LPLs in the bilayer (*e.g.* t = 300-320 *ns*, highlighted with a circle), these clusters are not long-lived.

### Mixed Bilayer Properties

The properties of the 20% LPLs bilayers were compared to those containing only phospholipids under equilibrium conditions. To increase sampling, larger bilayers were generated (Methods).

#### Bilayer Thickness

Bilayer thickness was calculated over the final 500 *ns* of each simulation using LiPyphilic. ^44–46^ Each leaflet was divided into a 10*×*10 grid and the bilayer thickness calculated in each bin as the distance between phosphorus atoms in the two leaflets. The bilayer thickness returned is the mean across all bins. The phospholipid-only bilayers were found to be slightly thicker than those containing LPLs: the bilayers containing only POPG, POPE, and POPA displayed a mean (*±* standard deviation) thickness of 3.917 *±* 0.025 *nm*, whereas the 20% LPLs bilayers displayed a thickness of 3.826 *±* 0.027 *nm*.

The partial densities of different groups within the systems were calculated over the *z*-axis to visualise these differences. Densities were calculated for lipid phosphorus atoms, acyl tails, glycerol groups, and water molecules over the final 500 *ns* of each simulation. Segmenting the *z*-axis into 100 bins, these distributions were similar across the two configurations (Fig. 10). The main difference is a subtle shift towards a thinner bilayer for the LPL-containing systems. For each moiety, there is a small shift in the distributions towards the centre of the bilayer (*z* = 0) for the 20% LPLs systems: the hydrophobic core is thinner and the distance between glycerol/phosphate groups is reduced on inclusion of LPLs. Furthermore, the hydroxyl groups of the LPLs are more hydrophilic than the ester groups that link the headgroup to the acyl tails. The effective size of the polar headgroup is thus increased in a lysophospholipid compared to its diacyl counterpart, which allows water to penetrate deeper into the bilayer than in a pure phospholipid membrane. ^17^

**Figure 10:**
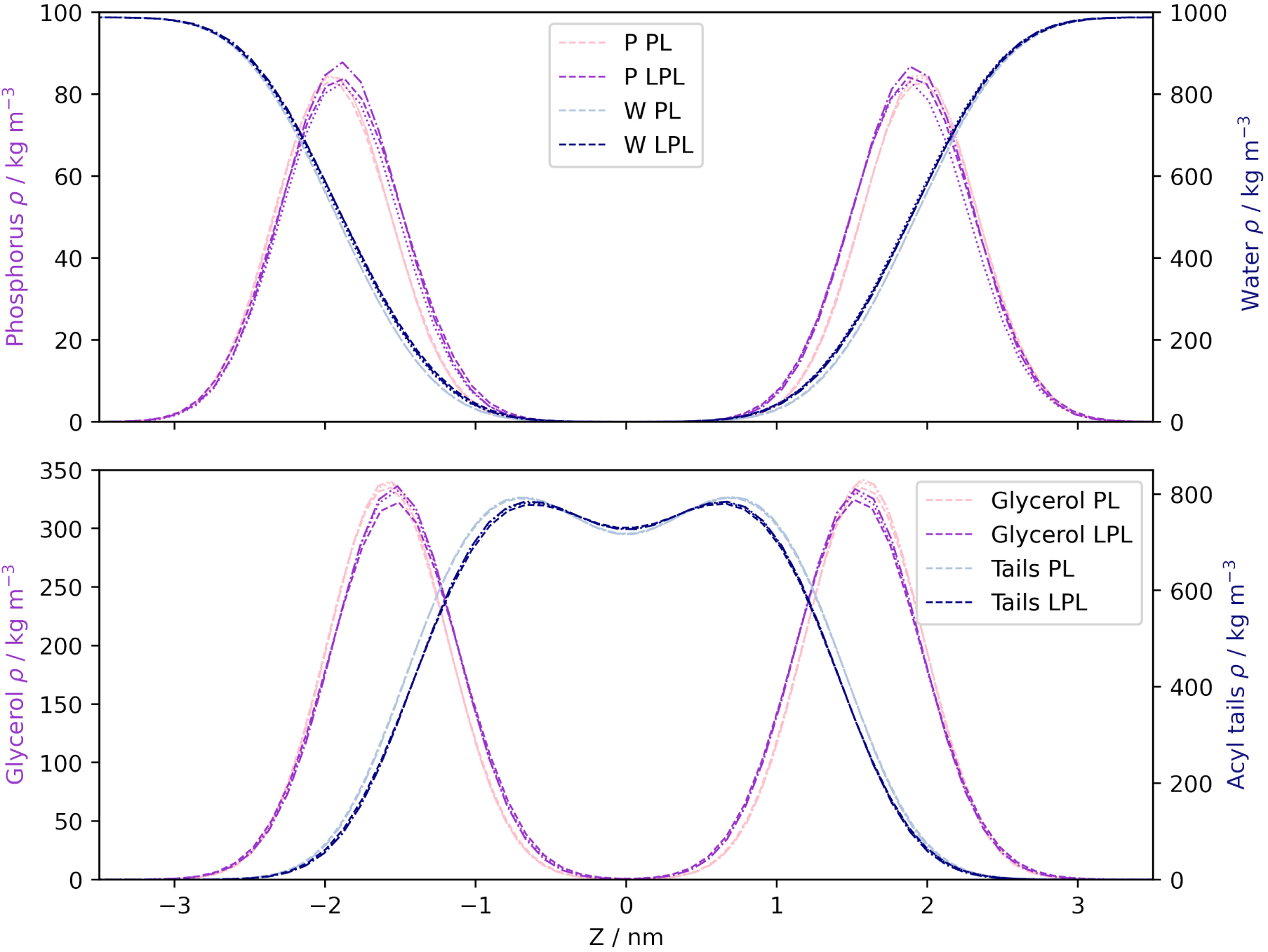
Partial densities of lipid moieties and water molecules in equilibrium simulations of the model bilayers. PL indicates phospholipid-only systems; LPL indicates 20% LPL systems. Top: Lipid phosphorus atoms and water molecules. Bottom: Lipid glycerol ester groups and lipid acyl tails. All partial densities are consistent with a subtle thinning of the bilayer on addition of LPLs.

#### Lipid Diffusion

Previous studies have shown that the addition of LPLs to egg phosphatidylcholine (16:0, 18:0 tails) bilayers can change membrane fluidity.^72^ Addition of lysoPC (16:0) significantly increased diffusion, while addition of lysoPE (16:0) slightly decreased diffusion.^72^ A combination of factors were cited, including changes in: van der Waals interactions; lipid packing; hydrogen bonding between phosphate groups and differing headgroups; and changes in bilayer thickness due to different tail lengths. ^72^ Owing to the complex mixture of lipids investigated here, it is difficult to predict how we would expect the diffusion of each lipid in the bilayer to change with composition.

The lateral diffusion constant for each lipid type in each bilayer was calculated over the final 500 *ns* of each simulation using the GROMACS utility msd. ^37,38^ These values are presented in Table 3. The average value for the phospholipid diffusion coefficients appears to decrease slightly when LPLs are included. However, the relatively large standard errors mean that the confidence in these values being representative of the true mean is low. We might expect a given lysophospholipid to display greater mobility compared to its diacyl counterpart, ^17^ e.g. lysoPE *versus* POPE. However, the standard error in these values again precludes any significant conclusions.

**Table 3:** Diffusion coefficients for the lipids in each bilayer system. Each value is presented as the mean *±* standard error of the mean across the three replicates of each system.

| | $D_{xy} / 1 \times 10^{-7} \text{cm}^2 \text{s}^{-1}$ | |
| --- | --- | --- |
|  | Phospholipid Only | 20% LPL Mix |
| **POPG** | $1.32 \pm 0.07$ | $1.20 \pm 0.25$ |
| **POPE** | $1.23 \pm 0.14$ | $1.16 \pm 0.15$ |
| **POPA** | $1.36 \pm 0.38$ | $1.24 \pm 0.25$ |
| **LysoPE**<sub>(18:1)</sub> | - | $1.29 \pm 0.30$ |
| **LysoPE**<sub>(16:0)</sub> | - | $1.33 \pm 0.69$ |
| **LysoPE**<sub>(19:0c)</sub> | - | $1.46 \pm 0.36$ |
| **LysoPG**<sub>(18:1)</sub> | - | $1.34 \pm 0.40$ |

#### Area Per Lipid

The Area Per Lipid (APL) was calculated over the final 500 *ns* of each trajectory for each system in two different ways. The average APL across all lipids in both leaflets was calculated by dividing the total area (twice the area of the *xy* plane) by the total number of lipids. The species-specific APL was then calculated using 2D Voronoi tessellation in LiPyphilic. ^46,73,74^ The calculated areas are presented in Table 4.

**Table 4:** Area per lipid (APL) for the lipids in each bilayer system. Each value is presented as the mean *±* standard error of the mean across the three replicates of each system. The first row indicates the average APL across all lipids, calculated using the total area of the two leaflets divided by the total number of lipids. Species-specific values were calculated using 2D Voronoi tessellation of atomic positions in LiPyphilic. The final column presents the percentage change between the phospholipid only system and the phospholipid-LPL system.

| | Area Per Lipid / nm<sup>2</sup> | | ΔAPL / % |
| --- | --- | --- | --- |
|  | Phospholipid Only | 20% LPL Mix |  |
| **All** | 0.6358 ± 0.0001 | 0.5977 ± 0.0001 | -5.99 |
| **POPG** | 0.6778 ± 0.0003 | 0.6471 ± 0.0003 | -4.53 |
| **POPE** | 0.6119 ± 0.0005 | 0.5797 ± 0.0004 | -5.26 |
| **POPA** | 0.6153 ± 0.0014 | 0.5761 ± 0.0011 | -6.37 |
| **LysoPE**<sub>(18:1)</sub> | - | 0.5639 ± 0.0011 | - |
| **LysoPE**<sub>(16:0)</sub> | - | 0.5542 ± 0.0011 | - |
| **LysoPE**<sub>(19:0c)</sub> | - | 0.5636 ± 0.0011 | - |
| **LysoPG**<sub>(18:1)</sub> | - | 0.6220 ± 0.0012 | - |

Three main observations were made from these values: the APL for a given lysophospholipid is smaller than that for its diacyl counterpart; the average APL across all lipids is reduced in bilayers containing the LPLs; and the APL for POPG, POPE, and POPA decreases on inclusion of LPLs. As the LPLs possess only one acyl tail, they occupy less space in the *xy* plane, leading to a reduced APL compared to the equivalent two-tailed lipid. Further to this, the LPLs appear to encourage closer packing of the phospholipids. If each phospholipid continued to occupy the same area as in a purely phospholipid bilayer, the reduced volume of the hydrophobic tails of the LPLs could lead to small vacuums in the regions around the LPLs tails. Thus, the lipids occupy less space in order to maintain the hydrophobic core of the bilayer.

#### Tail Order Parameters

The tail order parameter, *S_CD_*, is a measure of the orientational order of lipid acyl chain(s) in a bilayer. This can be calculated experimentally using quadrupolar splitting measured from deuterium NMR experiments, ^75,76^ and/or can be extracted from MD trajectories. ^77–81^ The order parameter for each carbon atom in the acyl chain is calculated as: ^75,76^

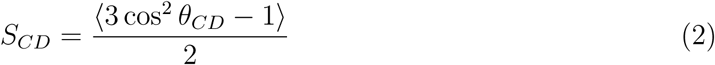

where *θ_CD_* is the angle between the bilayer normal (*z*) and the carbon-hydrogen/deuterium bond vector. Angular brackets indicate the ensemble average (molecular and temporal). Where the carbon has more than one bonded hydrogen, *S_CD_*may be calculated for each individual C-H bond, though through rotation these hydrogens are generally considered equivalent and the average C-H bond vector is used. A value of 0 indicates an isotropic ensemble with no overall preference in orientation, with positive and negative values indicating alignment with the bilayer normal or bilayer plane, respectively.

While there are several tools available to calculate order parameters,^82–85^ many rely on a united-atom approach which yields inaccurate results for unsaturated carbons. ^80^ Due to these inaccuracies, and difficulties in implementing existing all-atom tools, an in-house MDAnalysis script was used (available on Zenodo - see Supplementary Materials).

*S_CD_*was calculated separately for each phospholipid (POPG, POPE, POPA) acyl chain across all lipids of the same species over the final 100 *ns* of each replicate. All three phospholipids display a small decrease in order on inclusion of LPLs (Fig. 11). As discussed in the previous section, the packing of the lipids in the bilayer changes on addition of LPLs to maintain the hydrophobic core: the APL is reduced in order to maximise hydrophobic interactions between the acyl tails. However, due to the conical shape of LPLs there will still be an increased amount of free space within the core for the mixed bilayers compared to those containing only phospholipids. As a result, the phospholipid tails exhibit greater mobility and thus slightly reduced order parameters in the mixed bilayer systems.

**Figure 11:**
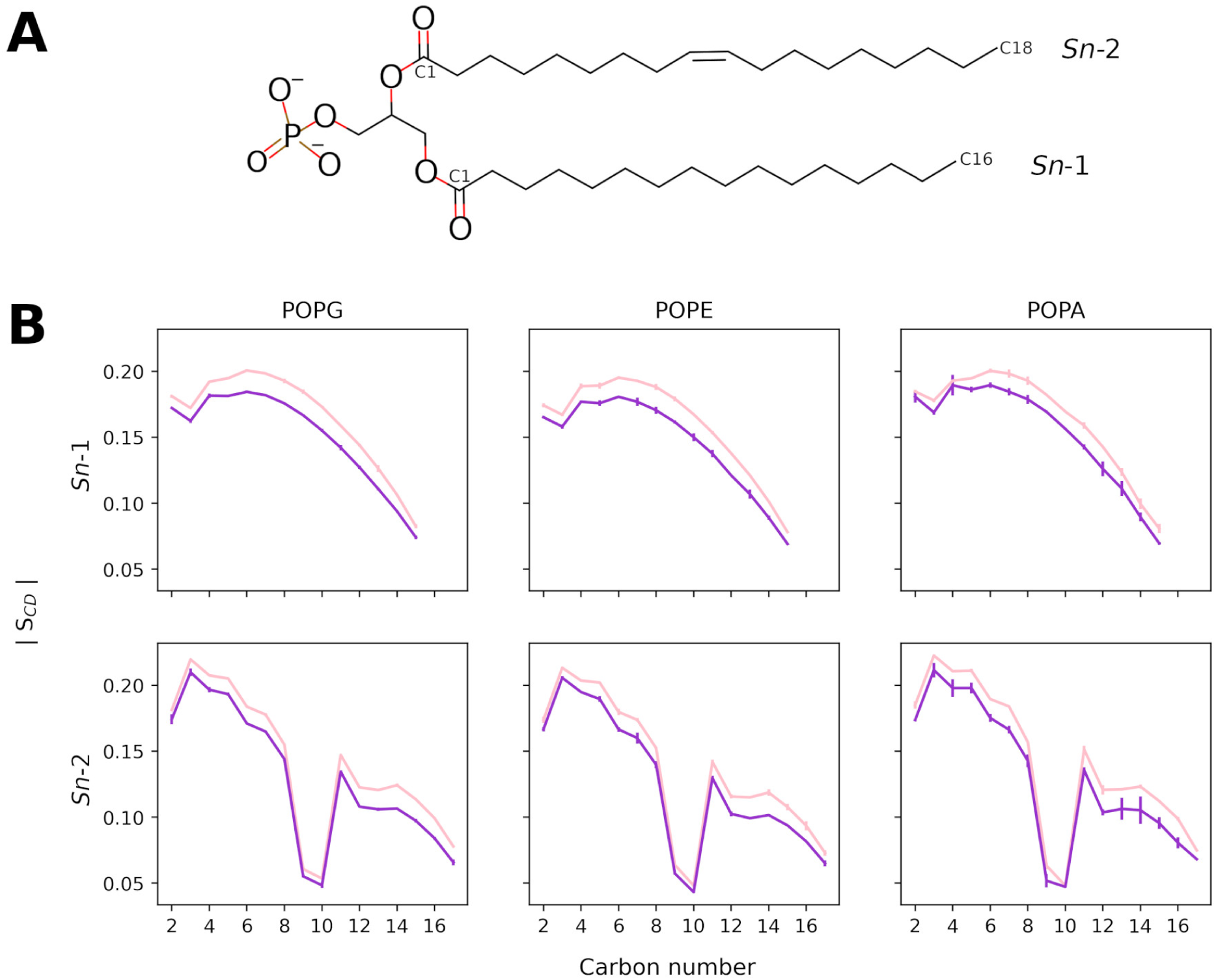
Comparison of tail order parameters for the phospholipids in different bilayer environments. **(A)** Structure of the phospholipid POPA with the *Sn*-1 and *Sn*-2 tails labelled, with start and end carbon atoms labelled on each chain. POPE and POPG share this tail structure. **(B)** Magnitude of the order parameter *S_CD_* for each carbon in each tail. Pink line indicates the average over replicates containing only phospholipids; purple line indicates 20% LPLs systems. Error bars show the standard deviation across the three replicates of each system. In all cases, the presence of LPLs reduces the order of the tails.

### Native protein embedded in Mixed Bilayer

We further tested the mixed membrane model by embedding the protein PglB in a pre-equilibrated bilayer. PglB is a native inner membrane protein in *C. jejuni* with oligosaccharyltransferase activity, needed for the transfer of glycans to acceptor proteins. This protein itself is glycosylated, with the acceptor sequon in the periplasmic domain. Three replicas of a system containing the mixed bilayer and one glycosylated PglB with its acceptor peptide and donor lipid-linked oligosaccharide were simulated for 1*µs* each.

#### Membrane Properties

Overall, the bilayer maintained its stability in all replicas, even though slight local curvature was induced around the protein. (Supplementary Materials, Fig. A2). Lipid tail order parameters computed for each phospholipid type were consistent among all replicas (Fig. 12A) and directly comparable with the order parameters reported for the mixed bilayer alone (Fig. 11).

**Figure 12:**
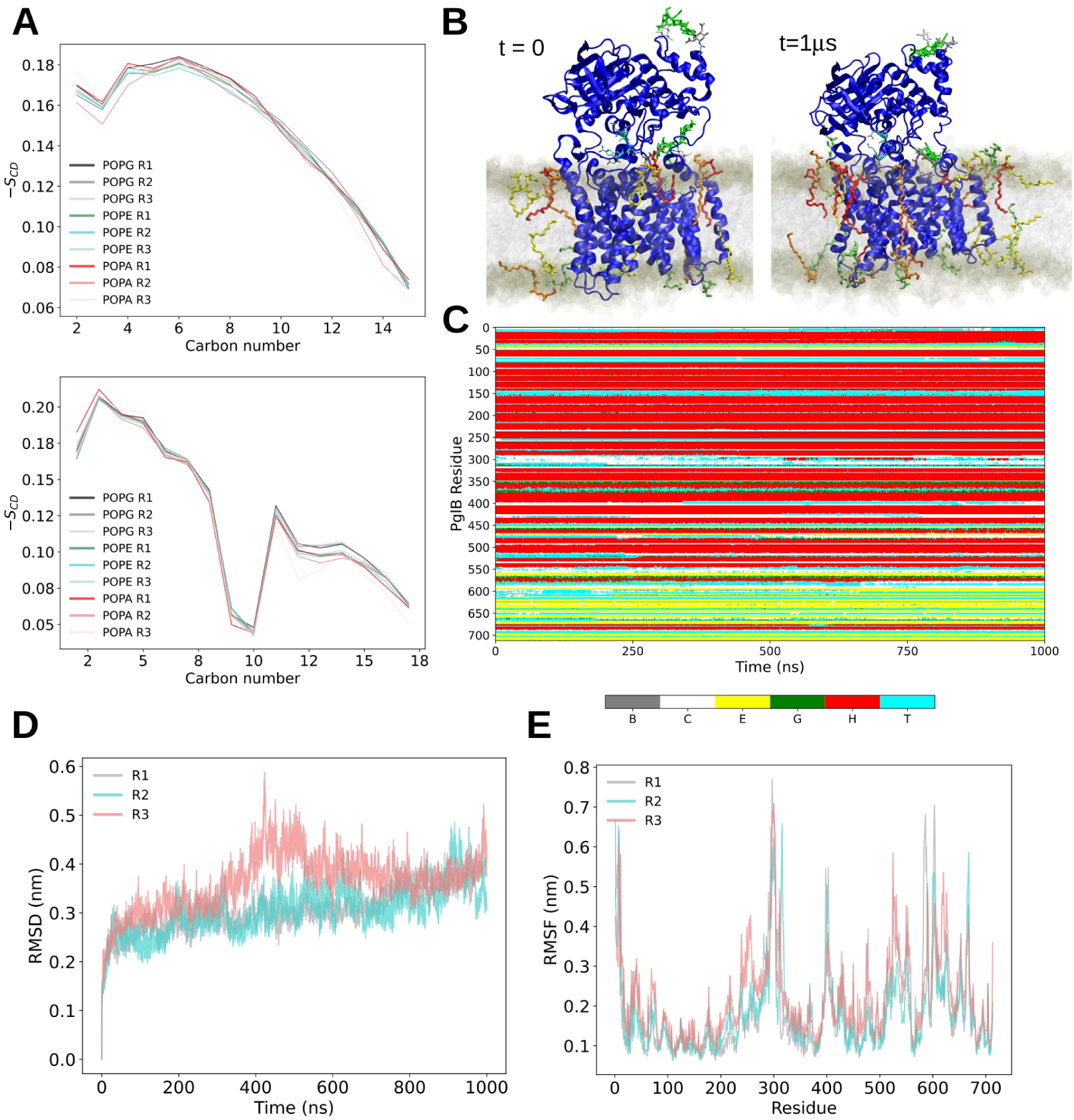
**(A)** Deuterium tail order parameters computed separately for all the phospholipids in each replica (R1-R3). **(B)** Zoomed-in side view of the system at the beginning and end of the production run for one example simulation (R1). The membrane is depicted as a translucent grey surface with phosphorus atoms of the lipids coloured in brown. PglB is shown as a blue cartoon representation. The acceptor sequon is shown with a cyan cartoon representation. Sugar components are shown by residue-based coloured sticks (white:, grey: green:). LPL that are in contact with PglB are depicted with residue-based coloured sticks inside the bilayer. (LysoPE_(18:1)_ : red, LysoPE_(16:0)_) **(C)** Secondary structure analysis for PglB over time in a representative replica (R1). STRIDE was used to compute the following secondary structure elements: isolated bridge (B, grey), coil (C, white), extended conformation (E, yellow), 3-10 helix (G, green), alpha-helix (H, red), turn (T, cyan). **(D)** Root Mean Square Deviation (RMSD) computed over the production run for all three replicas (R1-R3). **(E)** Root Mean Square Fluctuation (RMSF) of PglB computed on the production run for all three replicas.

For all replicas, the mean thickness values (*±* standard deviation) computed over the last 500 *ns* spanned values between 3.806-3.813 *±* 0.041 *nm*. These values are comparable with the ones previously mentioned for a bilayer-only system (3.826 *±* 0.027). The results show that the total average APL is consistent between the replicas (spanning values between 0.670-0.673 *±* 0.001 nm^2^, Table 5). The computed local APL showed slightly more variation between replicas, particularly so for the LPLs as shown in Table 5. Perhaps, the most noticeable cases are LysoPE_(18:1)_ and LysoPG_(18:1)_, which show a difference in APL of around 0.10 nm^2^ between replica 1 (R1) and 2-3. Towards the end of one of the trajectories, (R1) one LysoPG_(18:1)_ molecule moved such that its headgroup was well outside the plane of the headgroups of the other lipids, where it appeared to be stabilised by interactions with asparagine residues on the surface of PglB. (Supplementary Materials, Fig. A3). This event only occurred in one simulation and thus the significance is unclear.

**Table 5:**
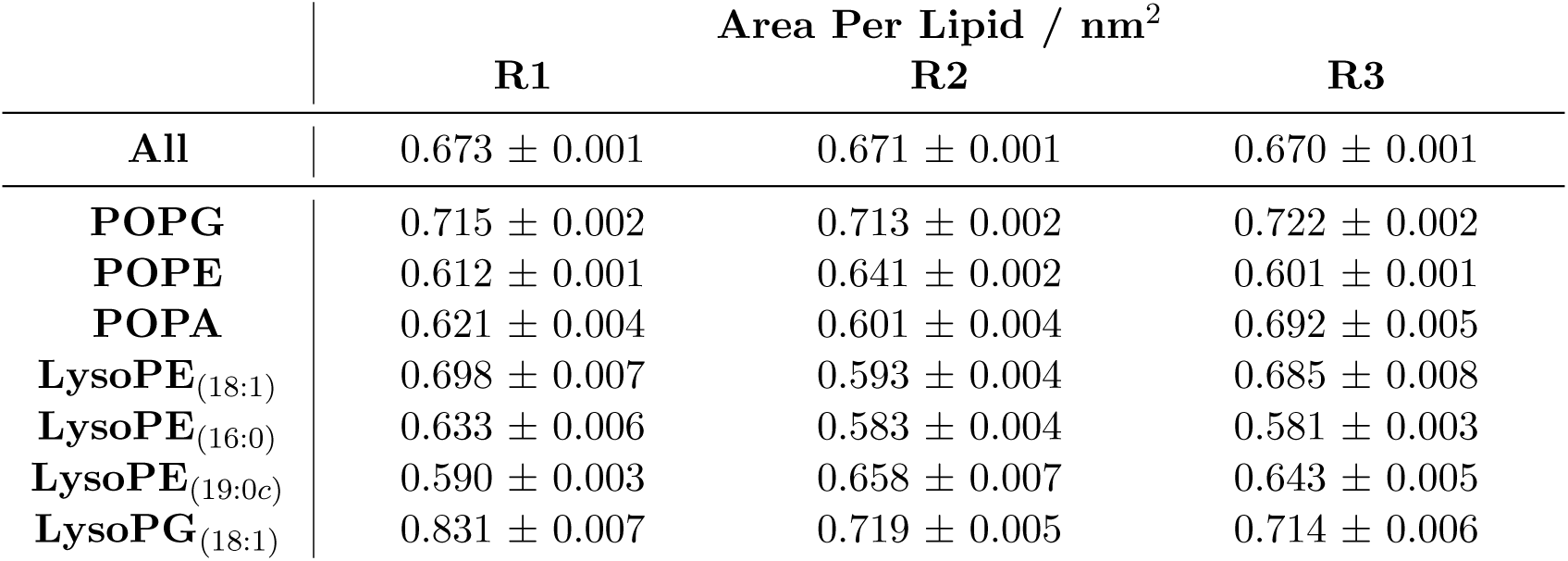
Area per lipid (APL) for the lipids in each protein-bilayer system. Each value is presented as the mean *±* standard error of the mean across the last 500 *ns* of each simulated system. Species-specific values were calculated using 2D Voronoi tessellation of phosphorus atomic positions of the lipids as implemented in fatslim. ^63^

#### Protein and Glycan Dynamics

Overall, PglB remained stably embedded inside the membrane for all replicas.(Supplementary Materials, Fig. A2). The secondary structure elements of the protein remained mainly stable along the simulation as shown in Fig. 12C. Some dynamical behaviour can be observed around the region spanning residues 290-300, a mainly unstructured region that is highly mobile and whose structural prediction, as reported in the AlphaFold model present in Uniprot, have low to very low confidence score (including a section with pLDDT *<* 50). The RMSD of the PglB backbone over time shows that the overall conformation of the protein is still changing slightly (particularly for replica R3), but in all cases it reached values around 0.4 *nm* by the end of the simulation. This could be rationalised by considering a local conformational change around the region binding the acceptor sequon peptide. In two replicas (R2 and R3) this peptide spontaneously dissociated from its binding site. The residue-based root mean square fluctuation was also computed and shown in Fig. 12E. All three replicas show the same qualitative profile indicating similar local dynamical behaviour. In the case of R3, although a new peak appears in the region spanning residues 250 - 270, that corresponds to regions spanning two distal transmembrane helices, overall, the RMSF profile is qualitatively comparable with the other simulated systems. Finally, we assessed the interactions of the glycans with the bilayer. Although PglB is glycosylated, this analysis was carried out on the carbohydrate elements of the glycosylated lipid donor, since it is the only molecule in this system with its glycan moiety proximal to the membrane. The sugar-lipid contacts were computed considering the lipids separately (see methods for more details). A dynamic behaviour was observed, where in all cases contacts could be established and lost in an unbiased way (Supplementary Materials, Fig. A4 and Table A9).

### Electroporation

Application of an electric field perpendicular to the bilayer has been explored extensively *via* MD as a method to induce pores^86–92^ - a process known as electroporation. The mechanism of electroporation begins with the formation of a water wire through the bilayer, favouring locations with local defects in the headgroup region. ^88,89^ The water wire is stabilised through interactions with lipid headgroups, which move into the hydrophobic core to further stabilise the water within the bilayer. ^89,90^ This allows the water wire to expand, resulting in pores in the membrane. In simulations this will often cause the rapid acceleration of water molecules into the channel and expansion of the pore in the bilayer plane. As the presence of varied lipid types results in a greater number of defects in the bilayer due to changes in the hydrophobic packing of the tails and hydrogen bonding between headgroups, we hypothesised that the presence of LPLs will allow electroporation at lower field strengths. To compare the integrity of the bilayers containing the lysophospholipids *versus* those without, we subjected these bilayers to an electric field normal to the membrane plane.

Equilibrated bilayers containing either 20% LPLs or exclusively phospholipids were subjected to a constant electric field along the *z*-axis, with field strengths from 0.1 to 0.2 *V nm^−^*^1^. The time taken for the bilayer to electroporate was then measured as the time at which the box *x* (equivalent to *y*) dimension increases to *>* 10% greater than the equilibrium box dimensions (Supplementary Materials, Fig. A5), i.e. the bilayer is expanding in the *xy* plane to accommodate a large pore that has formed in the membrane.

The electroporation times indicate that bilayers containing LPLs are more susceptible to electroporation than those containing only phospholipids (Table 6). Phospholipid-only bilayers were resistant to electroporation on the simulated timescales up to field strengths of 0.200 *V nm^−^*^1^, whereas bilayers containing LPLs were susceptible to electroporation at 0.150 *V nm^−^*^1^, and could not withstand field strengths greater than this. This increase in susceptibility to electroporation is consistent with the increase in tail disorder. Due to the stochastic nature of the membrane defects that lead to electroporation, the time taken for the membranes to electroporate is variable across replicates. The pores were found to close when the porated bilayers were subjected to equilibrium conditions. The time taken for the pores to close was measured as the time at which all water had retreated from the hydrophobic core. On average, these pores closed faster in the phospholipid-only systems (Table 6).

**Table 6:**
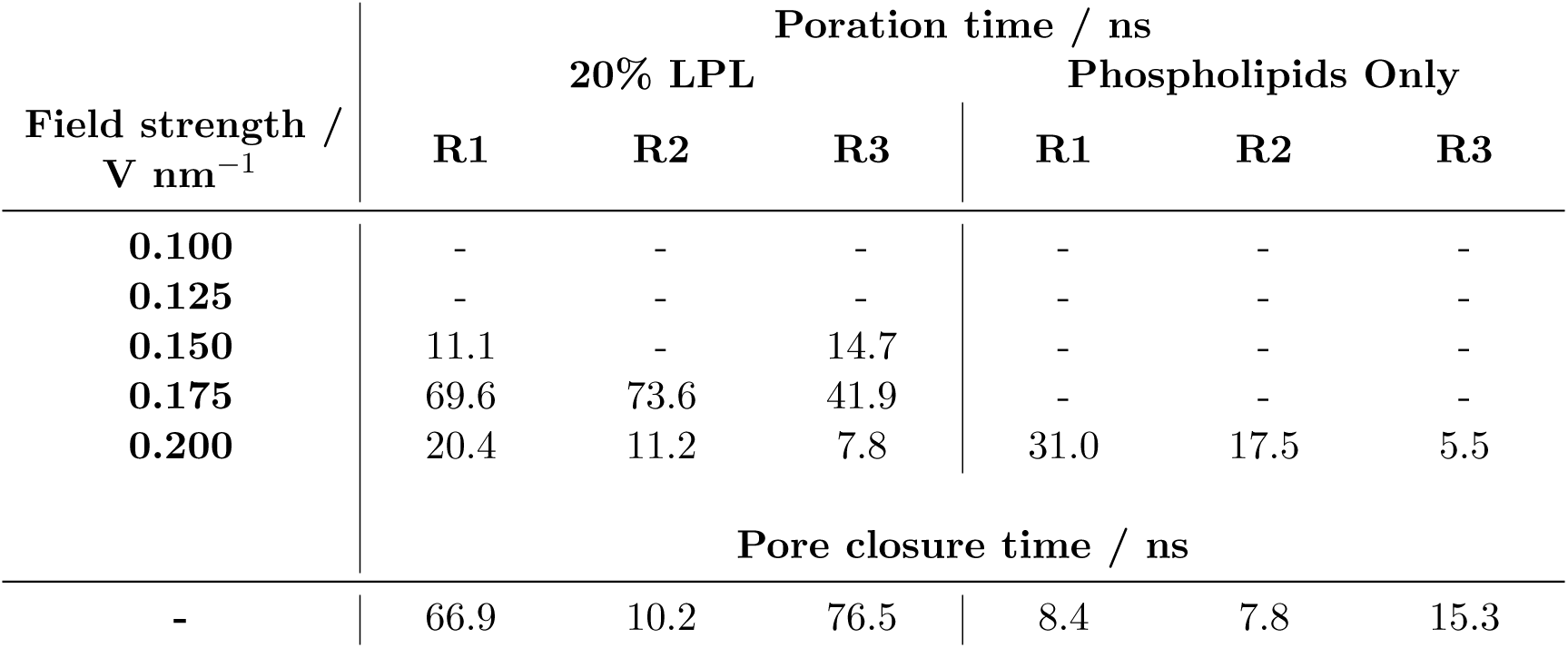
Electroporation times for the modelled bilayers under different field strengths and pore closer times under equilibrium conditions. Poration time measured as the time at which there is a large pore in the bilayer, resulting in the *x* dimension increasing by *>*10% compared to the equilibrium box dimensions. Pore closure time measured as the time at which all water had retreated from the hydrophobic core.

| Field strength /<br>$\text{V nm}^{-1}$ | Poration time / ns | | | | | |
| --- | --- | --- | --- | --- | --- | --- |
|  | 20% LPL |  |  | Phospholipids Only |  |  |
|  | R1 | R2 | R3 | R1 | R2 | R3 |
| **0.100** | - | - | - | - | - | - |
| **0.125** | - | - | - | - | - | - |
| **0.150** | 11.1 | - | 14.7 | - | - | - |
| **0.175** | 69.6 | 73.6 | 41.9 | - | - | - |
| **0.200** | 20.4 | 11.2 | 7.8 | 31.0 | 17.5 | 5.5 |
|  | Pore closure time / ns |  |  |  |  |  |
| - | 66.9 | 10.2 | 76.5 | 8.4 | 7.8 | 15.3 |

LysoPC has been shown experimentally to reduce electrical resistance and increase permeability of phosphatidylcholine bilayers. ^24,93^ The lysophospholipids in those bilayers are hypothesised to spontaneously form ion channels in the bilayer wherein two pairs of dimerised LPLs (each pair containing one lysophospholipid from each leaflet) stabilise a channel through the membrane (Fig. 14).^24^ The pores formed in the 20% LPLs bilayers appear to support this mechanism (Fig. 13C,D). Initial water wires form at positions where there are proximal LPLs in both leaflets, with the headgroups of these LPLs (and other nearby phospholipids) moving towards the bilayer centre to stabilise the pore.

**Figure 13:**
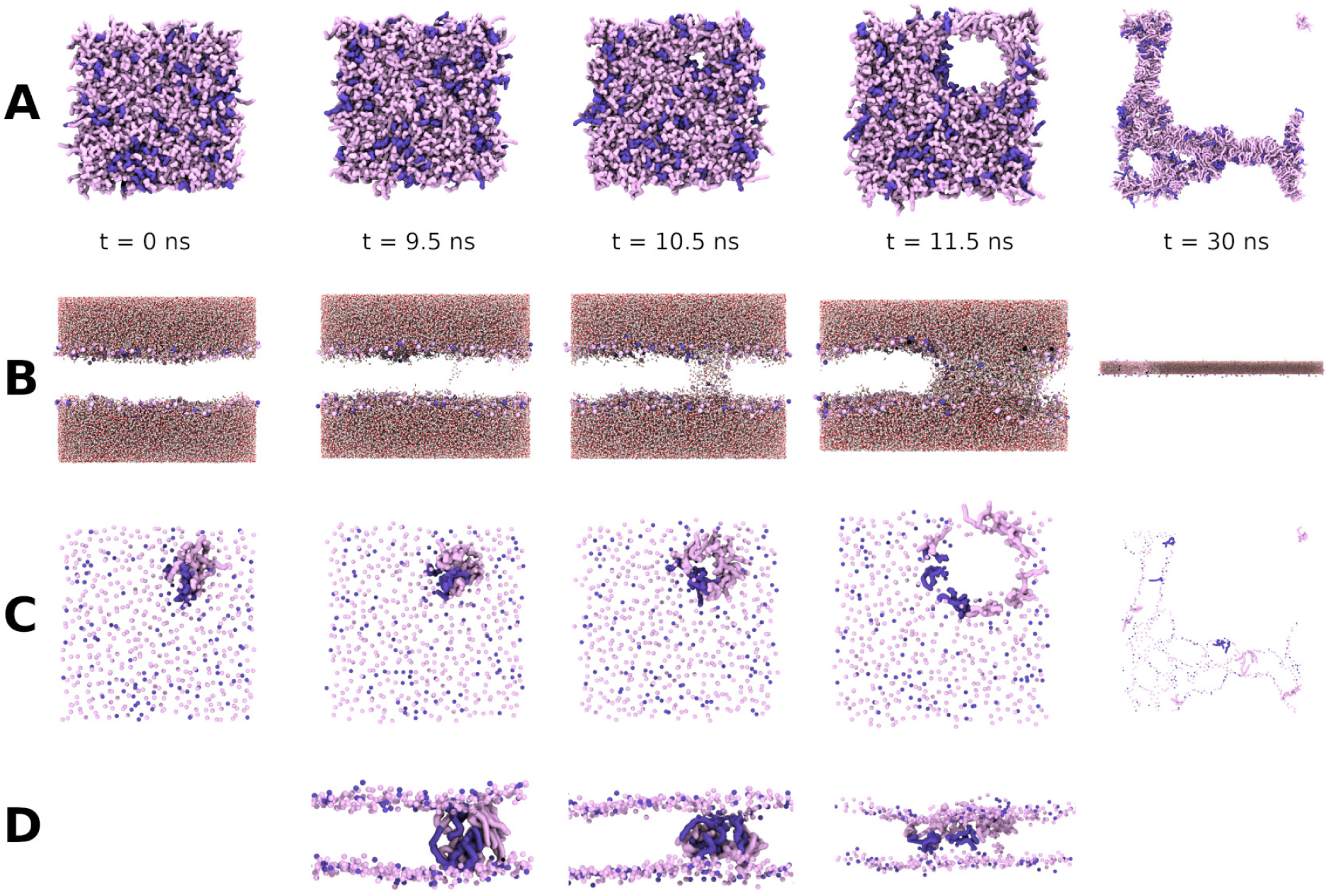
Electroporation of a 20% LPL bilayer (replicate 1, field strength 0.15 *V nm^−^*^1^). POPG, POPE, POPA in light pink, LPLs in purple. Water omitted in panels A, C, and D for clarity. **(A)** Top-down view of the bilayer at different stages of electroporation. **(B)** Side view of the bilayer at different stages of electroporation. Only phosphorus atoms of the lipids are shown for clarity. Water molecules shown as sticks coloured by element. **(C)** Top-down view of the bilayer showing lipid phosphorus atoms as spheres and lipids that line the initial pore shown in surface representation. **(D)** Side view of the bilayer showing lipid phosphorus atoms as spheres and lipids that line the initial pore shown in surface representation.

**Figure 14:**
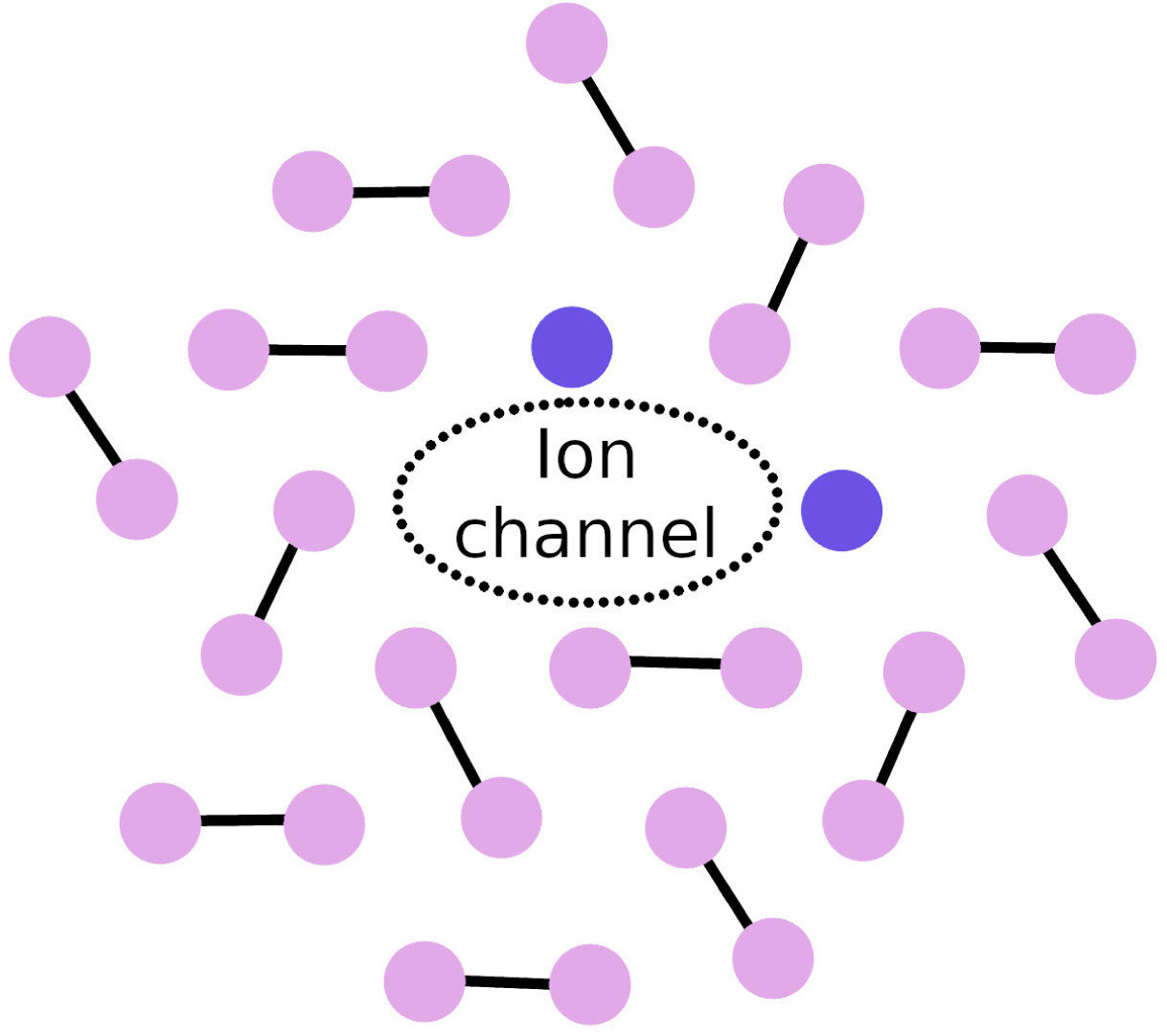
Schematic illustration of the cross-section of proposed ion channel formation in a phosphatidylcholine/lysophosphatidylcholine bilayer, adapted from Ref. 24. Phospholipids shown in pink, LPLs in purple. It is proposed that LPLs dimerise (one LPL from each leaflet), and pairs of dimers facilitate the formation of ion channels through the bilayer. We similarly observe pairs of LPLs at electroporation sites, suggesting that this mechanism may apply to both spontaneous pore formation and electroporation (Fig. 13).

## Limitations

It is useful to reflect on any potential limitations of the current study. Perhaps the most important, while we have qualitatively evaluated these models against general properties of this lipid family, the absence of quantitative data on the physical properties of these specific lipids in the literature has precluded quantitative assessment.

Our simulations in which phase behaviour was tested were somewhat limited by the system size and pressure coupling regimes. Small system sizes lead to difficulties in reproducing bulk properties as there is a requirement for periodic replication of the unit cell. Similarly, only the use of anisotropic coupling can allow the box to deform in such a way that accessible phases are not limited by box dimensions and enforcement of equal deformation in all dimensions. However, it can also lead to enormous deformation, to the point where the box collapses; this was observed in several simulations.

The electric field strengths utilised in the electroporation simulations are an order of magnitude greater than those employed in experimental studies, ^86^ as is usual for MD simulations given the much shorter timescales compared to laboratory experiments. The field strengths applied here are consistent with other MD studies. ^86,87,94,95^ Previous studies have also shown that the probability of observing poration increases with bilayer patch size. ^94^ While we attempted to generate bilayers of comparable size for these simulations, the phospholipid-only bilayers were slightly larger than their 20% LPLs counterparts (*x* dimensions of 13.5 *nm versus* ∼13.0 *nm*, respectively; ∼4% larger). Whether this small difference in size influenced the field strength required for electroporation was not investigated here. However, the phospholipid-only bilayers were the larger of the two configurations yet required greater field strengths to electroporate - the notion that LPLs reduce the integrity of the bilayer holds.

Finally, the complexity of the bilayers generated still fall short of the complete *C. jejuni* lipidome, which has been shown to contain more than 200 lipid species. ^9^ Here we have selected 4 lysophospholipids to model and our bilayers contain 7 lipid species in total; the tail and headgroup diversity *in vivo* is substantially greater than that modelled. Notable species that we have omitted include: lysophospholipids with myristoyl tails (C_14_), which have recently been highlighted as a novel virulence factor for *C. jejuni* ; ^96^ acylphospholipids and phospholipids containing cyclic moieties in their tails. ^9^ The latter species have greater acyl tail volume which can complement the lysophospholipids; including acylphospholipids and a greater proportion of lipids with cyclic moieties may be necessary to accommodate higher levels of LPLs^9^ in model membranes.

## Conclusions

In this work we have developed and tested atomistic models for four lysophospholipids found in the *C. jejuni* lipidome. We have shown these models to qualitatively match the expected behaviours of this family of lipids: micelles are formed at low lipid concentration, but at higher concentrations hexagonal, lamellar, and cubic phases may form. It was shown that solutions of a phospholipid-lysophospholipid mixture could self-assemble into bilayers on the sub-microsecond timescale. These bilayers were found to be slightly thinner than their phospholipid-only counterparts. While we did not observe any significant changes in lipid diffusion across the two bilayer compositions, the presence of LPLs was shown to reduce the APL and tail order parameters for each phospholipid species in the bilayer. We have shown that a glycosylated native inner membrane is conformationally stable within the mixed lipid bilayer and does not distort the local environment. Furthermore, these mixed membranes displayed increased susceptibility to electroporation compared to bilayers without LPLs.

## Acknowledgement

K.E.N. is supported by EPSRC (PhD Studentship Project Number: 2446840). A.F.B is funded by EP/V030779/1. S.K. is funded by EPSRC Grants EP/X035603, EP/V030779/1 and EP/Y008693/2. J.W.E. is funded by EP/Y008693/2. This work used the ARCHER2 UK National Supercomputing Service. The authors acknowledge the use of the IRIDIS High Performance Computing Facility, and associated support services at the University of Southampton, in the completion of this work.

## Supporting Information Available

Trajectories, run input files, custom analysis scripts, and topologies are available on Zenodo (DOI: 10.5281/zenodo.14833724). System contents and supplementary figures can be found in the Supplementary Information.

## Abbreviations

APL: Area Per Lipid
C*_W_*: Water weight fraction
CPP: Critical Packing Parameter
LLO: Lipid-Linked Oligosaccharide
LMPG: 1-myristoyl-2-hydroxy-sn-glycero-3-phospho-(1’-rac-glycerol)
LPLs: Lysophospholipids
MD: Molecular Dynamics
NMR: Nuclear Magnetic Resonance
PA: Phosphatidic acid
PC: Phosphatidylcholine
PE: Phosphatidylethanolamine
PG: Phosphatidylglycerol
PME: Particle Mesh Ewald
PMPE: 1-palmytoil-2-cis-9,10-methylenehexadecanoyl-phosphatidylethanolamine
POPA: 1-palmitoyl-2-oleoyl-sn-glycero-3-phosphatidic acid
POPE: 1-palmitoyl-2-oleoyl-sn-glycero-3-phosphoethanolamine
POPG: 1-palmitoyl-2-oleoyl-sn-glycero-3-phosphoglycerol
RMSD: Root Mean Square Deviation
RMSF: Root Mean Square Fluctuation

