## Supplementary Materials for "Atomistic modelling of lysophospholipids from the Campylobacter jejuni lipidome"

### Supplementary Material

#### Simulation System Contents

##### Self-assembly Simulations, Excess Water

Table A1: System contents for self-assembly simulations in excess water

| | **Small Single LPL systems** | | | | **Larger Single LPL systems (Anisotropic)** | | | |
|---|---|---|---|---|---|---|---|---|
| | **LysoPE$_{(18:1)}$** | **LysoPE$_{(16:0)}$** | **LysoPE$_{(19:0c)}$** | **LysoPG$_{(18:1)}$** | **LysoPE$_{(18:1)}$** | **LysoPE$_{(16:0)}$** | **LysoPE$_{(19:0c)}$** | **LysoPG$_{(18:1)}$** |
| **LysoPE$_{(18:1)}$** | 140 | 0 | 0 | 0 | 500 | 0 | 0 | 0 |
| **LysoPE$_{(16:0)}$** | 0 | 140 | 0 | 0 | 0 | 500 | 0 | 0 |
| **LysoPE$_{(19:0c)}$** | 0 | 0 | 140 | 0 | 0 | 0 | 500 | 0 |
| **LysoPG$_{(18:1)}$** | 0 | 0 | 0 | 140 | 0 | 0 | 0 | 500 |
| **Water** | 7,000 | 7,000 | 7,000 | 7,000 | 31,059 | 29,373 | 31,967 | 33,002 |
| **$K^+$** | 10 | 10 | 10 | 150 | 40 | 40 | 40 | 500 |
| **$Cl^-$** | 10 | 10 | 10 | 10 | 40 | 40 | 40 | 0 |
| **Initial box dimensions / nm** | 7.0×7.0×7.0 | | | | 10.9×10.9×10.9 | | | |

#### Self-assembly Simulations, $C_W$=0.4

Table A2: System contents for self-assembly simulations at $C_W$=0.4

| | Small Single LPL systems | | | | Larger Single LPL systems (Anisotropic) | | | |
|---|---|---|---|---|---|---|---|---|
| | **LysoPE**$_{(18:1)}$ | **LysoPE**$_{(16:0)}$ | **LysoPE**$_{(19:0c)}$ | **LysoPG**$_{(18:1)}$ | **LysoPE**$_{(18:1)}$ | **LysoPE**$_{(16:0)}$ | **LysoPE**$_{(19:0c)}$ | **LysoPG**$_{(18:1)}$ |
| **LysoPE**$_{(18:1)}$ | 140 | 0 | 0 | 0 | 500 | 0 | 0 | 0 |
| **LysoPE**$_{(16:0)}$ | 0 | 140 | 0 | 0 | 0 | 500 | 0 | 0 |
| **LysoPE**$_{(19:0c)}$ | 0 | 0 | 140 | 0 | 0 | 0 | 500 | 0 |
| **LysoPG**$_{(18:1)}$ | 0 | 0 | 0 | 140 | 0 | 0 | 0 | 500 |
| **Water** | 2,512 | 2,377 | 2,585 | 2,870 | 8,874 | 8,392 | 9,134 | 9,429 |
| $K^+$ | 2 | 2 | 2 | 142 | 20 | 20 | 20 | 500 |
| $Cl^-$ | 2 | 2 | 2 | 2 | 20 | 20 | 20 | 0 |
| **Initial box dimensions / nm** | 5.8×5.8×5.8 | | | | 9.9×9.9×9.9 | | | |

#### Self-assembly Simulations, $C_W$=0.1

Table A3: System contents for self-assembly simulations at $C_W$=0.1

| | Small Single LPL systems | | | | Larger Single LPL systems (Anisotropic) | | | |
|---|---|---|---|---|---|---|---|---|
| | **LysoPE$_{(18:1)}$** | **LysoPE$_{(16:0)}$** | **LysoPE$_{(19:0c)}$** | **LysoPG$_{(18:1)}$** | **LysoPE$_{(18:1)}$** | **LysoPE$_{(16:0)}$** | **LysoPE$_{(19:0c)}$** | **LysoPG$_{(18:1)}$** |
| **LysoPE$_{(18:1)}$** | 140 | 0 | 0 | 0 | 500 | 0 | 0 | 0 |
| **LysoPE$_{(16:0)}$** | 0 | 140 | 0 | 0 | 0 | 500 | 0 | 0 |
| **LysoPE$_{(19:0c)}$** | 0 | 0 | 140 | 0 | 0 | 0 | 500 | 0 |
| **LysoPG$_{(18:1)}$** | 0 | 0 | 0 | 140 | 0 | 0 | 0 | 500 |
| **Water** | 419 | 396 | 431 | 478 | 1,479 | 1,399 | 1,523 | 1572 |
| **$K^+$** | 1 | 1 | 1 | 141 | 5 | 5 | 5 | 500 |
| **$Cl^-$** | 1 | 1 | 1 | 1 | 5 | 5 | 5 | 0 |
| **Initial box dimensions / nm** | 5.4×5.4×5.4 | | | | 9.9×9.9×9.9 | | | |

#### Bilayer Self-assembly Simulations

Table A4: System contents for bilayer self-assembly simulations

| | **Bilayer Self-Assembly** | | | |
|---|---|---|---|---|
| | **POPG** | **POPE** | **POPA** | **20% LPL** |
| **LysoPE**$_{(18:1)}$ | 0 | 0 | 0 | 7 |
| **LysoPE**$_{(16:0)}$ | 0 | 0 | 0 | 7 |
| **LysoPE**$_{(19:0c)}$ | 0 | 0 | 0 | 7 |
| **LysoPG**$_{(18:1)}$ | 0 | 0 | 0 | 7 |
| **POPG** | 140 | 0 | 0 | 63 |
| **POPE** | 0 | 140 | 0 | 42 |
| **POPA** | 0 | 0 | 140 | 7 |
| **Water** | 7,000 | 7,000 | 7,000 | 7,000 |
| **K**$^{+}$ | 10 | 10 | 150 | 87 |
| **Cl**$^{-}$ | 10 | 10 | 10 | 10 |
| **Initial box dimensions / nm** | $7.2 \times 7.2 \times 7.2$ | | | |

Table A5: System contents for equilibrium simulation of bilayers with and without lysophospholipids

| | **20% LPL** | | | **Phospholipids Only** | | |
|---|---|---|---|---|---|---|
| | **R1** | **R2** | **R3** | **R1** | **R2** | **R3** |
| **LysoPE$_{(18:1)}$** | 63 | 63 | 63 | 0 | 0 | 0 |
| **LysoPE$_{(16:0)}$** | 63 | 63 | 63 | 0 | 0 | 0 |
| **LysoPE$_{(19:0c)}$** | 63 | 63 | 63 | 0 | 0 | 0 |
| **LysoPG$_{(18:1)}$** | 63 | 63 | 63 | 0 | 0 | 0 |
| **POPG** | 567 | 567 | 567 | 828 | 828 | 828 |
| **POPE** | 376 | 376 | 376 | 552 | 552 | 552 |
| **POPA** | 63 | 63 | 63 | 92 | 92 | 92 |
| **Water** | 103,808 | 104,583 | 104,663 | 71,713 | 71,691 | 71,756 |
| **$K^+$** | 1,145 | 1,147 | 1,147 | 1,113 | 1,114 | 1,113 |
| **$Cl^-$** | 452 | 454 | 454 | 193 | 194 | 193 |
| **Initial box dimensions / nm** | 20.9×20.9×10.5 | | | 21.1×21.1×8.9 | 21.0×21.0×9.0 | 21.1×21.1×8.9 |

#### Native protein embedded in Mixed Bilayer

Table A6: System contents for native protein in mixed bilayer. PglB protein was glycosylated in Asn534 (more details in main Method section). Pept refers to the acceptor sequon peptide; LLO refers to the glycosylated lipid donor.

| | **20% LPL, R1-R3** |
|---|---|
| **LysoPE**$_{(18:1)}$ | 22 |
| **LysoPE**$_{(16:0)}$ | 24 |
| **LysoPE**$_{(19:0c)}$ | 23 |
| **LysoPG**$_{(18:1)}$ | 24 |
| **POPG** | 211 |
| **POPE** | 144 |
| **POPA** | 21 |
| **LLO** | 1 |
| **PglB** | 1 |
| **Pept** | 1 |
| **MG** | 2 |
| **Water** | 61,969 |
| $\mathbf{K^+}$ | 420 |
| $\mathbf{Cl^-}$ | 169 |
| **Initial box dimensions / nm** | 12.6×12.6×15.8 |

#### Electroporation Simulations

Table A7: System contents for electric field simulation of bilayers with and without lysophospholipids

| | 20% LPL | | | Phospholipids Only | | |
|---|---|---|---|---|---|---|
| | **R1** | **R2** | **R3** | **R1** | **R2** | **R3** |
| **LysoPE$_{(18:1)}$** | 28 | 28 | 28 | 0 | 0 | 0 |
| **LysoPE$_{(16:0)}$** | 28 | 28 | 28 | 0 | 0 | 0 |
| **LysoPE$_{(19:0c)}$** | 28 | 28 | 28 | 0 | 0 | 0 |
| **LysoPG$_{(18:1)}$** | 28 | 28 | 28 | 0 | 0 | 0 |
| **POPG** | 252 | 252 | 252 | 324 | 324 | 324 |
| **POPE** | 168 | 168 | 168 | 216 | 216 | 216 |
| **POPA** | 28 | 28 | 28 | 36 | 36 | 36 |
| **Water** | 40,150 | 40,122 | 40,194 | 40,320 | 40,323 | 40,323 |
| **$K^+$** | 489 | 490 | 492 | 457 | 458 | 458 |
| **$Cl^-$** | 181 | 182 | 184 | 97 | 98 | 98 |
| **Initial box dimensions / nm** | 13.0×13.0×10.9 | 13.1×13.1×10.8 | 12.9×12.9×11.1 | 13.5×13.5×10.4 | 13.5×13.5×10.4 | 13.5×13.5×10.5 |

#### Simulation box collapse

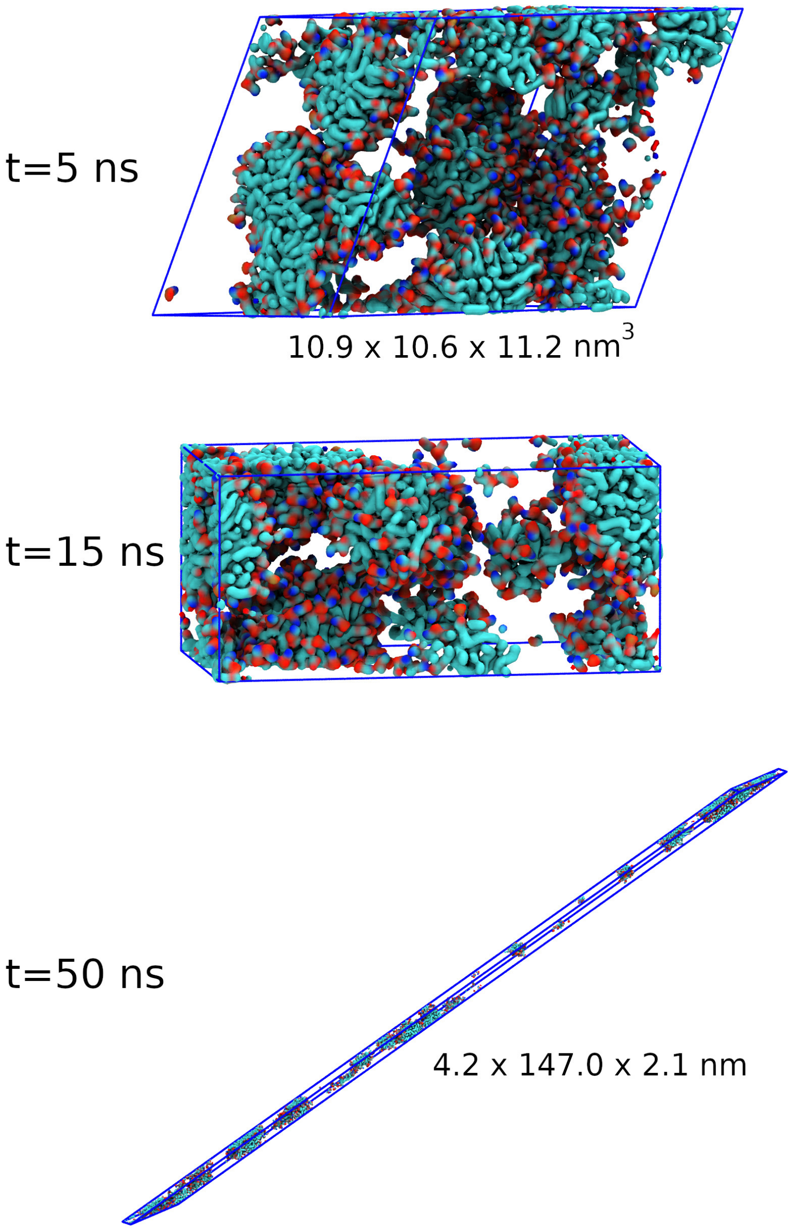


Figure A1: Collapse of a simulation box under anisotropic pressure coupling. Lipids shown as surfaces, coloured by element. Unit cell shown by blue lines. Simulation of lysoPE$_{(18:1)}$ (500 lipids, $C_W$=0.7) initially forms micelles. The unit cell deforms as the simulation progresses, resulting in box dimensions less than twice the electrostatics cut-off. The simulation crashes shortly after 50 ns.

### Depletion-enrichment indices

Table A8: Depletion-enrichment indices for lipids in the mixed bilayers. Values are presented as the mean ± standard deviation over the second half (final 500 ns) of each simulation. Calculated using the Python package LiPyphilic.[44–46,97]

| | | DEI | | | | | | |
|---|---|---|---|---|---|---|---|---|
| | | **POPG** | **POPE** | **POPA** | **lysoPE$_{(18:1)}$** | **lysoPE$_{(16:0)}$** | **lysoPE$_{(19:0c)}$** | **lysoPG$_{(18:1)}$** |
| **R1** | **POPG** | 0.95 ± 0.02 | 1.00 ± 0.02 | 0.92 ± 0.05 | 0.96 ± 0.05 | 1.06 ± 0.04 | 0.97 ± 0.06 | 0.95 ± 0.05 |
| | **POPE** | 1.05 ± 0.02 | 0.97 ± 0.04 | 1.11 ± 0.09 | 1.02 ± 0.08 | 0.95 ± 0.07 | 1.02 ± 0.07 | 1.10 ± 0.06 |
| | **POPA** | 0.96 ± 0.05 | 1.11 ± 0.10 | 0.90 ± 0.26 | 1.17 ± 0.28 | 0.90 ± 0.21 | 1.17 ± 0.22 | 0.93 ± 0.21 |
| | **lysoPE$_{(18:1)}$** | 1.02 ± 0.06 | 1.03 ± 0.09 | 1.19 ± 0.28 | 0.92 ± 0.21 | 1.18 ± 0.19 | 0.98 ± 0.19 | 1.11 ± 0.20 |
| | **lysoPE$_{(16:0)}$** | 1.12 ± 0.05 | 0.95 ± 0.07 | 0.90 ± 0.21 | 1.16 ± 0.20 | 0.8 ± 0.26 | 0.97 ± 0.22 | 0.93 ± 0.20 |
| | **lysoPE$_{(19:0c)}$** | 1.04 ± 0.06 | 1.04 ± 0.08 | 1.20 ± 0.23 | 0.99 ± 0.19 | 0.99 ± 0.23 | 1.09 ± 0.31 | 0.99 ± 0.17 |
| | **lysoPG$_{(18:1)}$** | 0.95 ± 0.05 | 1.04 ± 0.07 | 0.88 ± 0.20 | 1.04 ± 0.19 | 0.89 ± 0.20 | 0.93 ± 0.17 | 0.93 ± 0.28 |
| **R2** | **POPG** | 0.96 ± 0.02 | 0.99 ± 0.02 | 0.90 ± 0.05 | 0.98 ± 0.05 | 1.03 ± 0.06 | 0.97 ± 0.06 | 0.92 ± 0.05 |
| | **POPE** | 1.05 ± 0.02 | 0.98 ± 0.04 | 1.11 ± 0.06 | 1.01 ± 0.06 | 0.98 ± 0.08 | 1.00 ± 0.06 | 1.10 ± 0.07 |
| | **POPA** | 0.95 ± 0.05 | 1.11 ± 0.07 | 1.07 ± 0.39 | 1.08 ± 0.19 | 1.16 ± 0.25 | 1.07 ± 0.19 | 0.84 ± 0.24 |
| | **lysoPE$_{(18:1)}$** | 1.04 ± 0.05 | 1.02 ± 0.07 | 1.09 ± 0.18 | 0.83 ± 0.27 | 0.97 ± 0.21 | 1.15 ± 0.19 | 1.16 ± 0.26 |
| | **lysoPE$_{(16:0)}$** | 1.08 ± 0.06 | 0.97 ± 0.08 | 1.16 ± 0.24 | 0.96 ± 0.20 | 0.75 ± 0.25 | 0.95 ± 0.23 | 1.11 ± 0.19 |
| | **lysoPE$_{(19:0c)}$** | 1.04 ± 0.06 | 1.02 ± 0.07 | 1.09 ± 0.20 | 1.16 ± 0.19 | 0.96 ± 0.23 | 1.14 ± 0.26 | 1.06 ± 0.19 |
| | **lysoPG$_{(18:1)}$** | 0.93 ± 0.05 | 1.04 ± 0.07 | 0.80 ± 0.23 | 1.10 ± 0.25 | 1.06 ± 0.18 | 0.99 ± 0.17 | 0.95 ± 0.30 |
| **R3** | **POPG** | 0.95 ± 0.01 | 1.00 ± 0.02 | 0.90 ± 0.05 | 0.98 ± 0.05 | 1.01 ± 0.06 | 1.00 ± 0.05 | 0.95 ± 0.05 |
| | **POPE** | 1.06 ± 0.02 | 0.97 ± 0.04 | 1.13 ± 0.07 | 1.02 ± 0.07 | 0.95 ± 0.07 | 1.01 ± 0.06 | 1.05 ± 0.06 |
| | **POPA** | 0.95 ± 0.05 | 1.13 ± 0.08 | 0.86 ± 0.24 | 1.23 ± 0.22 | 1.08 ± 0.23 | 0.94 ± 0.17 | 1.04 ± 0.20 |
| | **lysoPE$_{(18:1)}$** | 1.04 ± 0.05 | 1.03 ± 0.08 | 1.24 ± 0.21 | 1.01 ± 0.32 | 0.99 ± 0.19 | 0.88 ± 0.22 | 1.06 ± 0.21 |
| | **lysoPE$_{(16:0)}$** | 1.07 ± 0.06 | 0.95 ± 0.08 | 1.09 ± 0.24 | 0.98 ± 0.18 | 1.04 ± 0.30 | 0.98 ± 0.22 | 1.14 ± 0.20 |
| | **lysoPE$_{(19:0c)}$** | 1.07 ± 0.06 | 1.02 ± 0.07 | 0.95 ± 0.17 | 0.88 ± 0.22 | 0.99 ± 0.22 | 1.01 ± 0.24 | 1.18 ± 0.18 |
| | **lysoPG$_{(18:1)}$** | 0.95 ± 0.05 | 1.0 ± 0.06 | 0.98 ± 0.19 | 1.0 ± 0.20 | 1.08 ± 0.19 | 1.1 ± 0.18 | 0.71 ± 0.30 |

### Membrane Protein

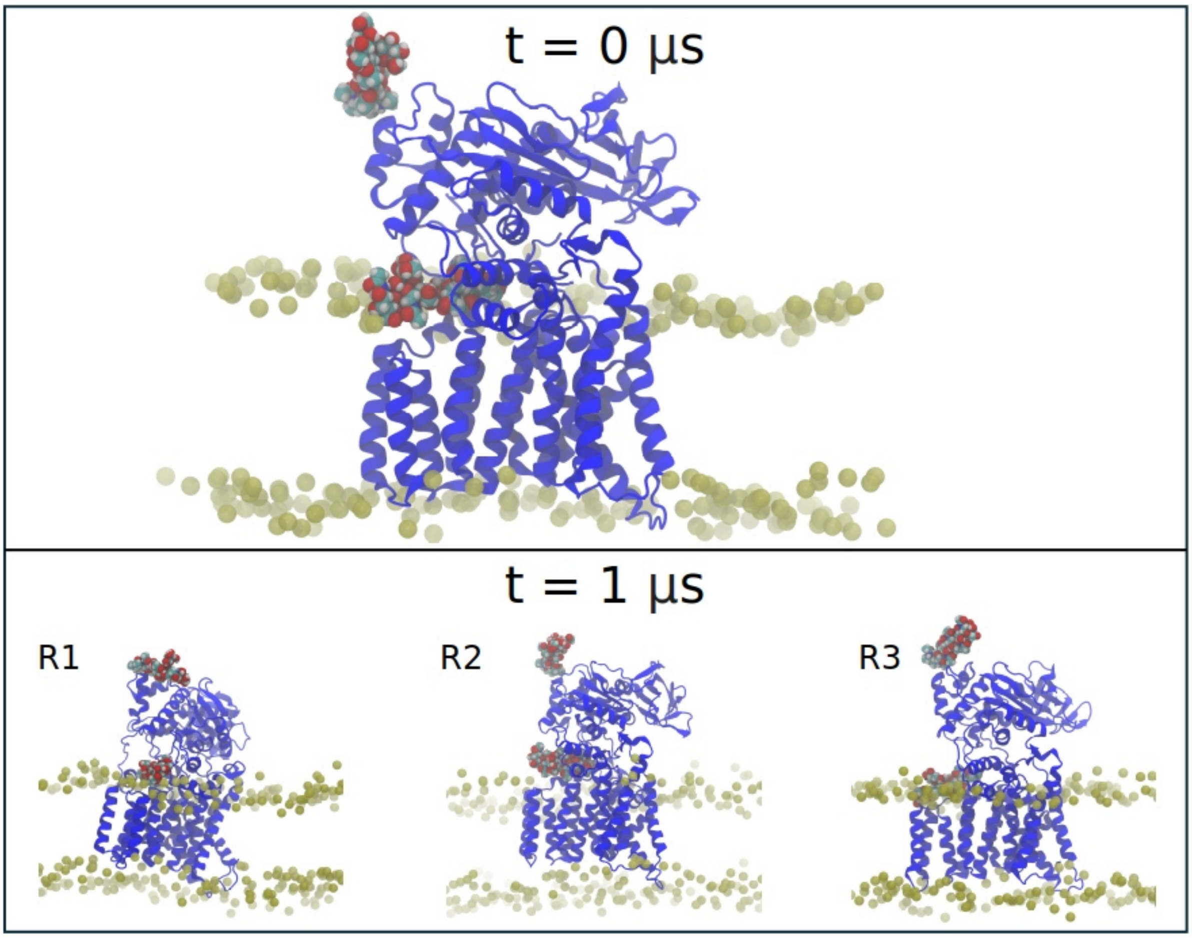


Figure A2: Zoomed-in sideviews of Initial and final snapshots of PglB/mixed bilayer for each of the replicas (R1-R3). PglB is coloured in blue cartoon representation, the glycans are depicted as coloured spheres and lipid phosphorus atoms are depicted as brown spheres.

Table A9: Average Sugar-lipid contacts over the whole trajectories

| | **Average Sugar-lipid contacts** | | |
|---|---|---|---|
| | **R1** | **R2** | **R3** |
| **POPG** | $0.416 \pm 0.008$ | $0.736 \pm 0.011$ | $1.021 \pm 0.011$ |
| **POPE** | $1.591 \pm 0.014$ | $0.386 \pm 0.009$ | $1.512 \pm 0.013$ |
| **POPA** | $0 \pm 0$ | $0.159 \pm 0.005$ | $0.266 \pm 0.006$ |
| **LysoPE**$_{(18:1)}$ | $0.0204 \pm 0.002$ | $0.137 \pm 0.005$ | $0.184 \pm 0.006$ |
| **LysoPE**$_{(16:0)}$ | $0 \pm 0$ | $0.042 \pm 0.003$ | $0 \pm 0$ |
| **LysoPE**$_{(19:0c)}$ | $0.002 \pm 0.001$ | $0.091 \pm 0.004$ | $0.014 \pm 0.002$ |
| **LysoPG**$_{(18:1)}$ | $0.019 \pm 0.002$ | $0.214 \pm 0.006$ | $0.021 \pm 0.002$ |

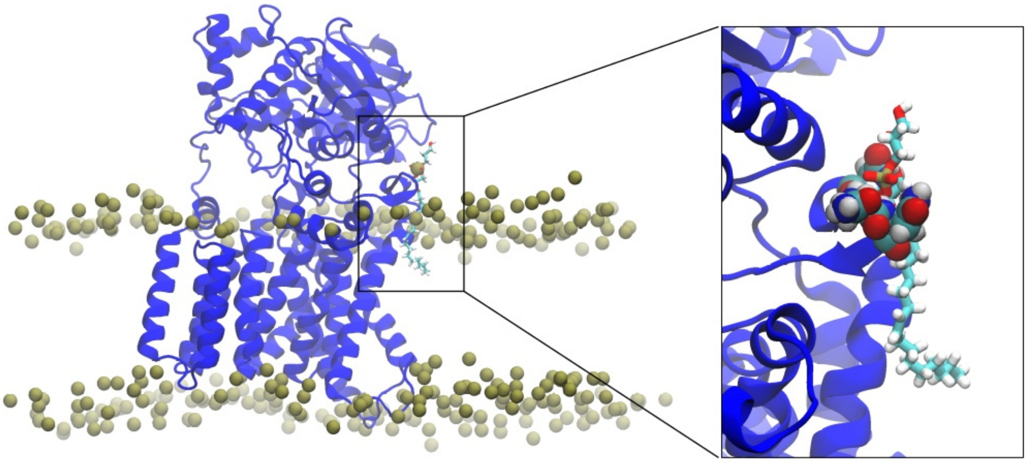

Figure A3: Final snapshot of R1 replica of PglB/mixed bilayer highlighting LysoPG$_{(18:1)}$ lipid (sticks) is located outside the bilayer alongside PglB (blue cartoon representation). Asparagine residues forming a 'cage' of interactions with the extracted LysoPG$_{(18:1)}$ are shown as spheres in the zoomed-in box. Lipid phosphorus atoms are depicted as brown spheres. Some phosphorus atoms in the immediate plane of vision in front of the protein have been omitted for clarity.

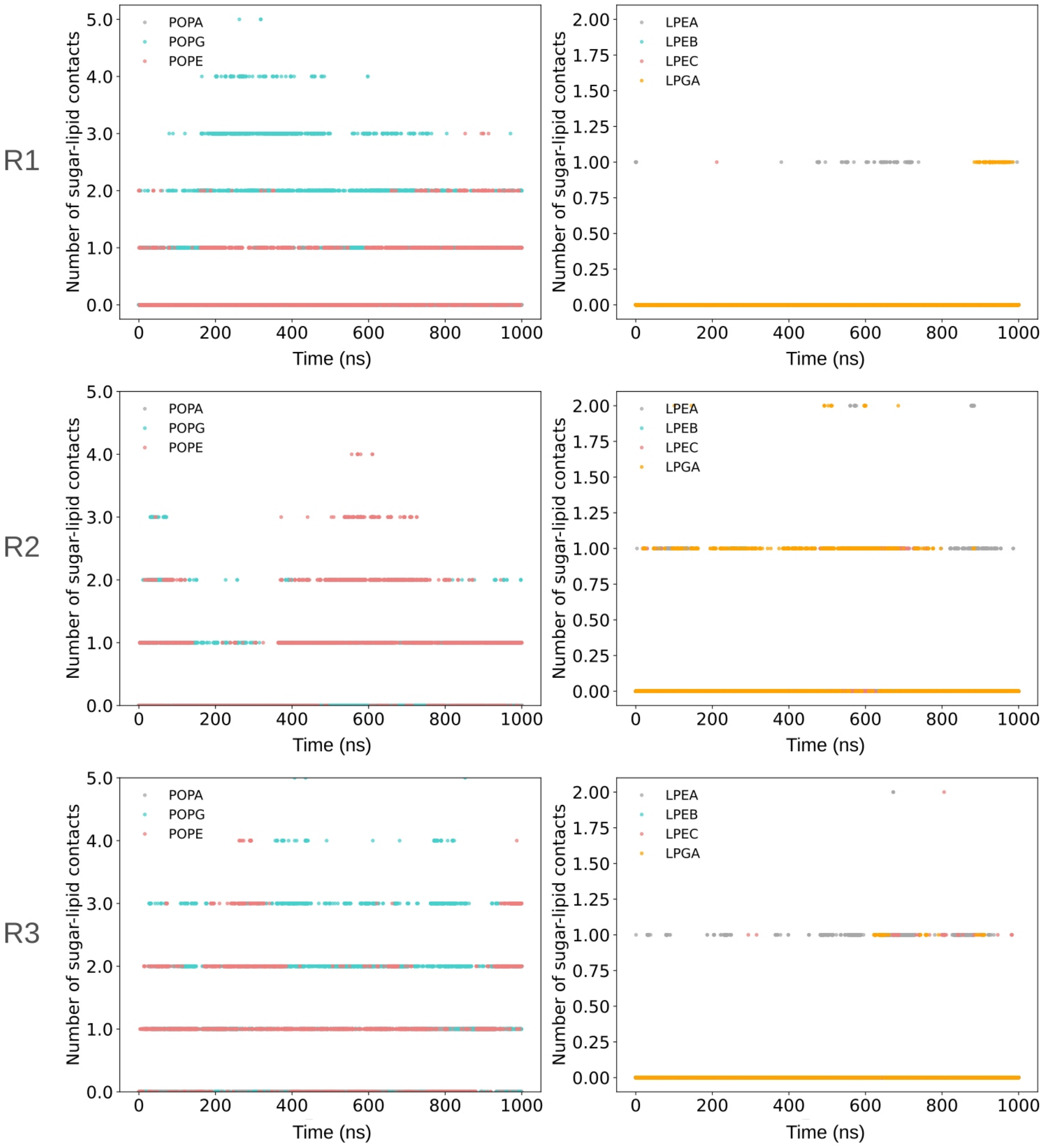


Figure A4: Contacts were computed between the sugar moieties of the LLO donor lipid and any lipid atom of the corresponding lipid type. Plots are separated for either phospholipids: POPA, POPG and POPE, or lysophopholipids: LPEA: LysoPE$_{(18:1)}$, LPEB: LysoPE$_{(16:0)}$ , LPEC: LysoPE$_{(19:0c)}$ , LPGA: LysoPG$_{(18:1)}$

### Electroporation Time

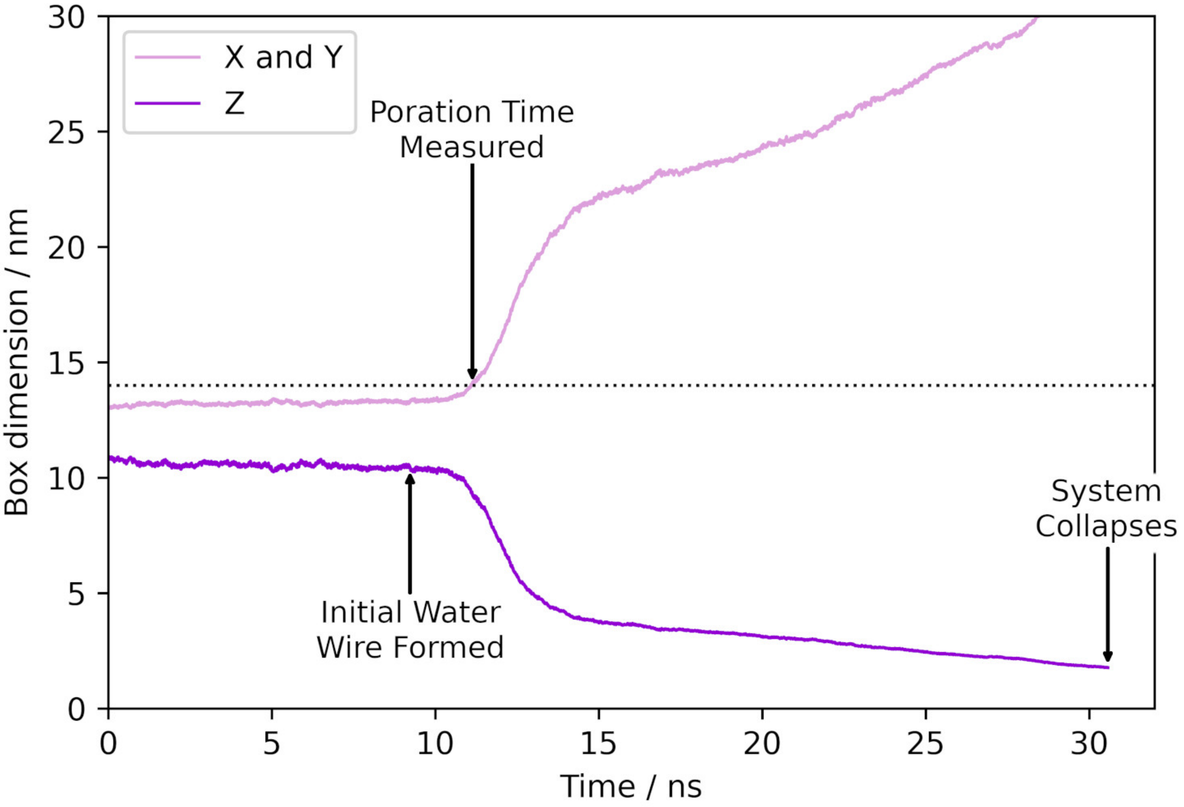


Figure A5: Box dimensions during electroporation of a mixed bilayer (R1, field strength 0.15 V $nm^{-1}$). The formation of the water wire has a negligible effect on the box dimensions, but as lipid headgroups move into the hydrophobic core to stabilise the water channel the bilayer expands in the $xy$-plane. The poration time was measured as the point at which the $x$ (and $y$) box dimension increased to >10% above the equilibrium box dimension; at this point, a substantial water channel has formed in the bilayer.
